# Tuning the double lipidation of salmon calcitonin to introduce a pore-like membrane translocation mechanism

**DOI:** 10.1101/2023.07.13.548826

**Authors:** Philip M. Lund, Kasper Kristensen, Nanna W. Larsen, Astrid Knuhtsen, Morten B. Hansen, Claudia U. Hjørringgaard, Anne Z. Eriksen, Andrew J. Urquhart, Kim I. Mortensen, Jens B. Simonsen, Thomas L. Andresen, Jannik B. Larsen

**Affiliations:** Center for Intestinal Absorption and Transport of Biopharmaceuticals, Technical University of Denmark, Lyngby, Denmark; DTU Health Tech, Department of Health Technology, Technical University of Denmark, 2800 Kgs. Lyngby, Denmark

**Keywords:** Lipidated peptide, membrane transport, membrane permeation, fluorescence microscopy, salmon calcitonin, Giant Unilamellar Vesicles

## Abstract

A widespread strategy to increase the transport of therapeutic peptides across cellular membranes has been to attach lipid moieties to the peptide backbone (lipidation) to enhance their intrinsic membrane interaction. Efforts *in vitro* and *in vivo* investigating the correlation between lipidation characteristics and peptide membrane translocation efficiency have traditionally relied on end-point read-out assays and trial-and-error-based optimization strategies. Consequently, the molecular details of how therapeutic peptide lipidation affects it’s membrane permeation and translocation mechanisms remain unresolved. Here we employed salmon calcitonin as a model therapeutic peptide and synthesized nine double lipidated analogs with varying lipid chain lengths. We used single giant unilamellar vesicle (GUV) calcein influx time-lapse fluorescence microscopy to determine how tuning the lipidation length can lead to an All-or-None GUV filling mechanism, indicative of a peptide mediated pore formation. Finally, we used a GUVs-containing-inner-GUVs assay to demonstrate that only peptide analogs capable of inducing pore formation show efficient membrane translocation. Our data provided the first mechanistic details on how therapeutic peptide lipidation affects their membrane perturbation mechanism and demonstrated that fine-tuning lipidation parameters could induce an intrinsic pore-forming capability. These insights and the microscopy based workflow introduced for investigating structure-function relations could be pivotal for optimizing future peptide design strategies.

**Graphical Abstract:** 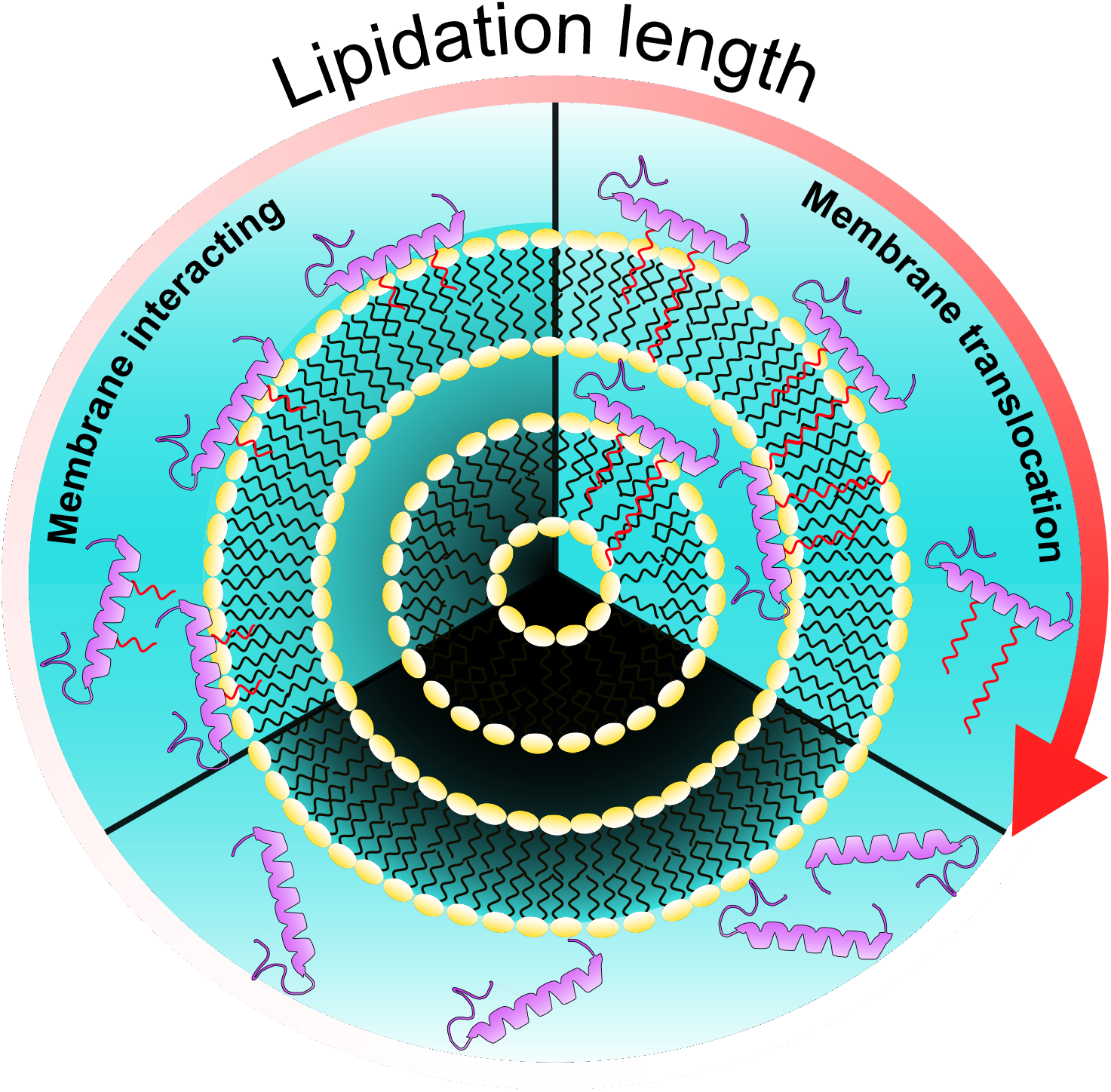

**Highlights:** - Lipidating the therapeutic peptide salmon calcitonin alters its biophysical characteristics, including oligomer size, hydrophobicity and membrane activity.

- Fluorescent microscopy of single GUVs enables the determination of peptide mediated reporter dye influx behavior as either graded or All-or-None, which is coupled to either smaller membrane perturbations or peptide pore formation.

- Modulating the number of hydrocarbons constituting the lipidation moieties determines the membrane permeation mechanism.

- By increasing the lipid chain length lipidated of salmon calcitonin goes from displaying smaller membrane perturbations to a peptide pore formation mechanism.

- Effective membrane translocation of lipidated salmon calcitonin requires a peptide mediated pore forming mechanism.

## Introduction

The size and hydrophilic nature of peptides most often prevent them from diffusing passively through cell membranes and cell-cell junctions [1]. Thus, one of the remaining challenges in delivering therapeutic peptides, especially through oral administration, is increasing the transport efficiency across the epithelial cell barrier [2–6]. A traditionally pursued strategy has been to covalently modify therapeutic peptides with lipid groups (lipidation) hereby increasing their intrinsic ability to interact with and cross the epithelial cell membranes [7,8]. This approach has been investigated for a plethora of therapeutic peptides pursued for oral delivery including insulin [9,10], for the treatment of type 1 and 2 diabetes [11]; glucagon-like peptide-2 [12], for the treatment of inflammatory bowel diseases and short bowel syndrome [13]; and salmon calcitonin (sCal) [14–17] for the treatment of various bone-related disorders, including osteoporosis, Paget’s disease, and hypercalcemia [18–22]. The majority of these studies focus on single-lipidated peptides, where the peptide performance is quantified using *in vitro* or *in vivo* experiments providing end-point measurements of how lipidation affected cross barrier transport. Thus, little is known about the molecular details of how therapeutic peptides translocate across membranes and how lipidation affects the membrane transport mechanism.

Detailed studies on membrane permeation have traditionally focused on membrane active peptides [23–27]. The membrane permeation mechanism of these peptides are conventionally determined via bulk leakage experiments based on large unilamellar vesicles (LUVs), including classical calcein leakage setups [12,16,26,28–31], ANTS/DPX requenching assays [26,31–33], or more recently fluorescent lifetime measurements of calcein release [24,34–36]. Based on the final integrity of the LUVs and the resulting leakage profile two mechanistic terms have been defined: All-or-None (AoN) and graded [31,37]. AoN describes a peptide displaying a strong permeation ability, which allows for full equilibration between the solution inside and outside the LUV and is thus associated with membrane pore formation. The graded mechanism is described to be based on numerous weak and short-lived peptide induced membrane perturbations leading to a partial exchange of solutions across the membrane.

A common feature of the used release assays is, that they provide a flexible high-throughput platform enabling a comparison of the averaged membrane permeation between different peptides. Nevertheless some of these assays are limited by being bulk readouts of many LUVs simultaneously, making it impossible to determine, if a recorded partial response in e.g. calcein release originates from all LUVs giving a partial response, or only some LUVs giving a full response and others showing no response. Thus the ensemble averaging in these assays can make assigning molecular mechanism difficult. Other bulk assays such as the ANTS/DPX assay circumvent this issue by first performing leakage experiments with LUVs containing the dye-quencher pair ANTS/DPX, followed by an additional around of DPX quencher, which can delineate the leakage mechanism, however these assays are limited by being end-point measurements, hampering the access to kinetic information [38,39]. To circumvent these potential issues investigators have implemented setups relying on imaging single giant unilamellar vesicles (GUVs) using time-lapse fluorescence microscopy [23,27,40–42]. Performing a spatiotemporal analysis on single fluorescently labeled GUVs confined in a solution containing a dye allows for direct observation of the dye filling kinetics into individual GUVs, which enables direct temporal delineation between AoN and graded mechanisms. This facilitates the assignment of the molecular mechanism underlying the peptide mediated membrane perturbations at the single-vesicle level. Wheaten *et al.* [43] further developed the single GUV assay by producing and observing GUVs-containing-inner-GUVs (GiGs), providing the opportunity to quantify indirectly the membrane translocation of label-free cell-penetrating peptides. This allowed them to move beyond the classical assessment of membrane activity and directly ascertain how the ability to interact with and permeate membranes was linked to cross-membrane translocation. However, none of the sophisticated imaging-based assays, have yet been applied for studying lipidated peptides, constituting one of the main reasons why we still lack molecular detail on how lipidation site and length affect membrane translocation of therapeutic peptides.

Here we used sCal as a model therapeutic peptide to study how the length of double lipidations affects its membrane permeation potency and mechanism. Nine lipidated variants of sCal were synthesized using the cross-combination of alkyl chains with the length of 4, 8 and 12 carbons. We found a complex relation between lipidation length, the peptide oligomer size and hydrophobicity, however, bulk calcein leakage data allowed us to classify the sCal variants into three groups with varying membrane activity. A sCal variant from each membrane activity group and the native sCal were further investigated in a fluorescence GUV microscopy assay to determine the underlying permeation mechanisms. Our results demonstrated that minor changes in lipid length lead to significant variation in the membrane permeation mechanism. A graded mechanism was found for short lipidations, whereas longer lipidations showed a time-dependent shift from a graded to an AoN mechanism, suggesting a pore-forming ability of these sCal variants. Evaluating the four selected sCal variants in the GiGs assay revealed translocation capabilities for only the sCal variants displaying an AoN mechanism, proposing pore formation as a prerequisite for efficient membrane translocation. Overall, our findings provide mechanistic insights on how lipidation affect peptide translocation across membranes. Such fundamental information could potentially provide rational design strategies for future efforts in peptide lipidation research.

## Materials and Methods

### Materials

All standard and orthogonally protected N_α_-Fmoc amino acids were purchased from Iris-Biotech (Marktredwitz, Germany). Pd(PPh_3_)_4_ was purchased from Alfa Aesar (Haverhill, MA, USA). All other reagents applied in peptide synthesis were purchased from Sigma Aldrich(St. Louis, MO, USA). Sodium dihydrogen phosphate, sodium hydroxide, hydrochloride acid (HCl), sodium chloride (NaCl), calcium chloride (CaCl_2_), magnesium chloride (MgCl_2_), tablets for phosphate-buffered saline (PBS), sucrose, D-(+)-glucose, nile red, pyrene, thioflavin T, *tert*-butanol, calcein, tetramethylrhodamine isothiocyanate–Dextran-4.4kDa (Dex), bovine serum albumin (BSA), phosphorus and gallium standards for ICP, and Triton X-100 were purchased from Sigma-Aldrich (St. Louis, MO, USA). Dimethyl sulfoxide (DMSO) and chloroform were purchased from VWR Chemicals (Radnor, PA, USA). 1-Palmitoyl-2-oleoyl-*sn*-glycero-3-phosphocholine (POPC) was purchased from Avanti Polar Lipids (Alabaster, AL, USA). Atto-655-1,2-Dioleoyl-sn-glycero-3-phosphoethanolamine (Atto655-DOPE) was purchased from Atto-Tec (Siegen, Germany). Black 96-well plates were purchased from Nunc, Thermo Fisher Scientific (Waltham, MA, USA). Slurry for preparing Sepharose CL-4B columns was purchased from GE Healthcare (Little Chalfont, UK). Econo-Column glass chromatography columns were purchased from Bio-Rad (Hercules, CA, USA). Protein LoBind tubes were purchased from Eppendorf (Hamburg, Germany). µ-Slide 8 well glass bottoms were purchased from Ibidi USA inc. (Fitchburg, WI, USA). All buffers and aqueous solutions were prepared using Milli-Q water (18.2 MΩ cm, prepared using a Milli-Q Academic, Millipore, Burlington, MA, USA).

### sCal variants synthesis and purification

Peptides were synthesized on a Tentagel S RAM amide resin (0.24 mmol/g) (Rapp-Polymer) using standard microwave Fmoc-synthesis on a Biotage Alstra peptide synthesizer. Specially synthesized amino acids (N_4_, N_8_ or N_12_; see supporting information) were incorporated using standard synthesis, followed by incorporation of the orthogonally protected Fmoc-L-Glu(OAll)-OH. After final automated synthesis, the batches were split in three and for each batch the OAll was removed and the respective amine chains were coupled as specified (see supporting information). Final cleavage was performed in a cleavage cocktail of 94% TFA, 2.5% EDT, 2.5% H_2_O and 1% triisopropylsilane, and the resin was stirred in the cocktail for 2.5 hours at room temperature before evaporation, Et_2_O-precipitation and lyophilization. The disulfide was formed using a stock solution 0.06M I_2_ in MeOH, which was quenched using ascorbic acid and the resulting peptide was lyophilized before purification using standard reverse-phase HPLC purification. See full experimental in the supplementary information.

### Dynamic Light Scattering (DLS)

100 µL solutions containing 100 µM peptide in phosphate buffer (10 mM phosphate, 100 mM NaCl, 1 mM CaCl_2_, 1 mM MgCl_2_, pH 6.7) were prepared in 1.5 mL Protein LoBind tubes. The solutions were centrifuged for 1 minute at 3300*g* using a MiniSpin centrifuge (Eppendorf) to precipitate impurities interfering with the DLS recording. 85 µL of the centrifuged solutions were transferred to plastic micro UV cuvettes (Brand, Wertheim, Germany). The cuvettes were incubated for at least 1 hour at 37 °C. The samples were then evaluated by DLS using a Zetasizer Nano ZS (Malvern, Worcestershire, UK) heated to 37 °C. The acquired correlation curves were treated using an intensity-based size distribution analysis. The peptide peaks were identified by comparing to the size distribution of a blank phosphate buffer sample.

### Nile red measurement

98 µL solutions with 100 µM peptide in phosphate buffer were transferred to a black 96-well plate. 2 µL of 1.5 mM nile red solution in DMSO was transferred to each well to a final nile red concentration of 30 µM. The solutions in the wells were mixed by pipette. The plates were incubated ∼1-2 hours at 37 °C before transfer to a Spark multimode microplate reader (Tecan, Männedorf, Switzerland) heated to 37 °C. The nile red fluorescence emission intensity of the samples at 633 nm was then measured using an excitation wavelength of 550 nm.

### Calcein release assay

The liposome calcein release is an established technique to determine membrane permeation[16,30,44]. LUVs consisting of POPC were loaded with self-quenched calcein. To prepare the LUVs, lipid powder was first dissolved and mixed in *tert*-butanol/water (9:1). The samples were then plunge-frozen in liquid nitrogen and lyophilized overnight (ScanVac CoolSafe lyophilizer, LaboGene, Allerød, Denmark). The lipids were hydrated with 1 mL of calcein solution (60 mM calcein, 10 mM phosphate, pH 6.7) to a lipid concentration of 50 mM. The samples were gently vortexed every 5 minutes for a total period of 30 minutes and subsequently subject to five freeze-thaw cycles by alternate placement in a liquid nitrogen bath and a 70 °C water bath. Next, the samples were extruded 21 times through a 100 nm polycarbonate filter (Whatman, GE Healthcare) using a mini-extruder (Avanti Polar Lipids) to form the LUVs. The LUVs were separated from free calcein using a Sepharose CL-4B size-exclusion chromatography column using phosphate buffer as the eluent (column dimensions: 1.5×20 cm, flow rate: 1 mL/min). To avoid osmotic strain on the membranes of the LUVs, the osmolality of the phosphate buffer (207 ± 1 mOsmol/kg) matched the osmolality of the calcein solution (211 ± 1 mOsmol/kg). After separation, the LUVs were transferred to an Amicon Ultra-4 100 kDa centrifugal filter unit (Merck, Darmstadt, Germany) and concentrated by centrifuging at 2000*g*. The phosphorus concentration of the concentrated samples was determined using inductively coupled plasma mass spectrometry (ICP-MS). For this purpose, the samples were diluted ×5100 in 2% HCl, 10 ppb gallium aqueous solution before performing the ICP-MS recording using an iCAP Q ICP-MS system (Thermo Fisher Scientific) operated in kinetic energy discrimination mode with gallium as the internal standard. The phosphorus concentration was determined by comparing it to a set of phosphorus standard samples in the range 25-100 ppb. The phospholipid concentration was determined by subtracting the background of the phosphate buffer. The lipid concentration of the concentrated samples was typically in the range of 11-14 mM. The size of the LUVs was assessed by DLS using a Zetasizer Nano ZS operated at 25 °C. The acquired correlation curves were treated using the cumulant analysis. The Z-average size and polydispersity index of the LUVs were 115 ± 2 nm and 0.06 ± 0.01, respectively. To perform the calcein release assay, the peptide and LUV solutions were heated to 37 °C. The peptides and LUVs were mixed directly in black 96-well plates. The final measured peptide concentrations were 100 µM, 30 µM, 10 µM, 3 µM, 1 µM, 0.3 µM or 0.1 µM, the final lipid concentration was 50 µM, and the final volume was 150 µL. The plates were incubated for 30 minutes at 37 °C. The calcein fluorescence emission intensity of the samples at 514 nm was then measured using an excitation wavelength of 491 nm. The percentage calcein release was calculated using equation 1.

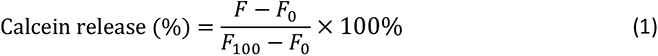

where *F* is the calcein fluorescence emission intensity of the given sample, *F*_0_ is the calcein fluorescence emission intensity of a non-treated sample, and *F*_100_ is the calcein fluorescence emission intensity of a sample with complete calcein release, obtained by incubating the LUVs with 0.5% Triton X-100 for 30 minutes at 37 °C.

### Preparation of Giant Unilamellar Vesicle

The GUVs were produced via electroformation using an in-house procedure adopted from previously reported protocols [23,43,45]. For POPC GUVs a 1 mM lipid stock solution in chloroform consisting of POPC:Atto655-DOPE in the mol% ratio 99.5:0.5 was mixed and stored at -20 °C until use. 20 μL of the lipid solution was added on two clean platinum rods, which were attached on a custom teflon support. Both the teflon support and rods were kept in vacuum for at least 1 hour ensuring effective chloroform evaporation. To produce the GUVs, the rods were next fully immersed into a 400 μL solution containing 226 mM sucrose and attached to a function generator (Digimess HUC6500 FG100, Stenson, England), producing an AC field between the two rods. To produce high-quality POPC GUVs, a series of six settings were applied: 1) 10 Hz and 0.2 V for 5 minutes, 2) 10 Hz and 0.5 V for 5 minutes, 3) 10 Hz and 1.0 V for 10 minutes, 4) 10 Hz and 1.5 V for 15 minutes, 5) 10 Hz and 2.0 V for 45 minutes and finally 6) 4 Hz and 2.0 V for 30 minutes. All voltages used were constant sine-wave shaped. Applied settings produced a mixed population of both single GUVs and GiGs. Assuming that all lipids on the rods formed GUVs, final lipid concentration was calculated to be 50 μM in the 400 μL sucrose solution. Following electroformation the sucrose solution containing newly formed GUVs was transferred to a clean glass vial and wrapped with aluminum foil to protect from light. The GUVs were stored at 4 °C and used within 48 hours.

### Spinning disk microscopy imaging of GUVs

Recording of GUVs was performed using a Nikon Ti2, Yokogawa CSU-W1 spinning disc confocal microscope equipped with a high numerical aperture 60× oil immersion objective and a Photometrics Prime 95B sCMOS detector. A 226 mM glucose solution with an osmolality of 228 mOsmol/kg was prepared with similar osmolality to the sucrose solution (235 mOsmol/kg) applied for GUV preparation, ensuring minimal osmotic strain on the GUVs. Next BSA was applied to passivate Ibidi micro-slide 8 wells by adding 300 μL 1 g/L BSA in PBS to each well for a minimum of 30 minutes before washing each well 8 times with 300 μL of the glucose solution. To perform GUV experiments, 150 μL glucose solution with 50 μM calcein was mixed in the well with 25 μL GUV solution and incubated for 5 minutes allowing the GUVs to settle. The difference in molar mass between the glucose solution added in the wells and sucrose solution used to form the GUVs, enable the GUVs to sediment on the bottom of the wells. Lipid concentration of the GUV solution, which was calculated to be 50 μM, provided a final concentration of 3.85 μM in the wells. A 150 μL solution containing 50 μM calcein and 325 μM peptide was prepared and carefully loaded into the well, resulting in final peptide concentration of 150 μM and a calcein concentration of 46 μM. Addition of the peptide solution fills the well to 100% capacity and ensures proper mixing. Subsequently, after adding the peptide solution, the GUVs were imaged for a minimum of 30 minutes, but maximum for 45 minutes, with a temporal resolution of 15 seconds. For experiments with TRITC-dextran-4kDa, the dye was added in the same amounts and solutions as calcein. The images were acquired by alternating between exciting calcein, TRITC-dextran-4kDa, and Atto655-DOPE using 488 nm, 560 nm and 638 nm diode laser lines, respectively. The calcein emission was detected through a 520/28 Brightline HC filter set, the TRITC-dextran-4kDa emission was detected through 600/50 ET bandpass filter set, and Atto655-DOPE emission was detected through 700/75 ET bandpass filter set.

### GUV image treatment and analysis

Recorded GUV images were analyzed with an in-house made python script. A flow chart (Fig. S1A) and an extensive description is found in the supplementary information. In brief, each image of the recorded series was flattened to eliminate illumination profile artifacts (Fig. S1B). GUVs were then located based on the Atto655 fluorescent signal of the membrane channel using Scikit-image [46], and linked using Trackpy [47]. The GUV membrane channel was applied to create a mask allowing for extraction of both calcein and TRITC-dextran-4kDa mean GUV lumen intensities both inside the GUV and outside the GUV representing the mean local background intensity (Fig. S1D). For extracting the calcein and TRITC-dextran-4kDa localized inside the GUV, the membrane mask was shrunk uniformly to define a pixel area localized inside the GUV. For extracting the local background a donut shaped pixel area beyond the GUV membrane mask was implemented. Each extracted GUV was manually accessed for quality unilamellarity. Single GUV traces lasting less than 20 frames were excluded to ensure a sufficient amount of time points for analysis. A 5 frames rolling average was implemented on the calcein and TRITC-dextran-4kDa GUV lumen intensities and background intensities to reduce noise. Percent filling for each time point (Percent filled *t_n_*) of the single GUV was calculated using equation 2.

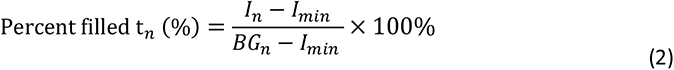

where *I_n_* is the mean GUV lumen intensity for the n’th time point, *I_min_* is the lowest intensity in the trace and *BG_n_* the local background for the n’th time point. The percent filling kinetic traces were manually fitted using SciPy [48]. To ensure high quality slope values from the fitting, selected traces consisting of less than five time points were not included in the slope data evaluation. Images were sectioned and edited for brightness and contrast in ImageJ. The recorded GUV population was manually assessed for GiGs data.

### Data presentation and statistical analysis

Matplotlib [49] was used for plotting data. SciPy and NumPy [50] were used for linear regression and numeric operations. Statistical analysis of distribution data was preformed using Cramér-von Mises’s (CVM) test to account for the non-Gaussian nature and unequal variance of the data.

## Results

### Synthesis design and biophysical characterization of double lipidated sCal variants

To identify a suitable strategy for lipidating sCal, we considered the sequence and structure of the peptide (Fig. 1A and 1B). sCal obtains a nearly random coil conformation in water, but in more structure-promoting solvents or in the presence of sodium dodecyl sulfate micelles, the central region of the peptide (approximately residues 8-22) forms an amphipathic α-helix, while the C-terminal residues remain mobile and folds back onto the core of the peptide [51–53]. Based on this structural information, we synthesized a library of nine sCal analogues with two saturated hydrocarbon chains attached. One of the chains was attached at the Q20 position to enhance the amphipathicity of the central helix, while the other chain was attached at the N26 position to increase the total lipophilicity of the peptide and potentially also promote a structure-stabilizing interaction with the Q20 position (Fig. 1A and 1B, Helical wheel representation was generated using HeliQuest [54]). Lipidated glutamine and asparagine were used in the peptide synthesis (Fig. 1C). Lipids with chain lengths of 4, 8, and 12 carbons were attached on each site producing the symmetrical and asymmetrical cross combinations of all three chain lengths. We refer to the peptides as CxCy, where Cx represents the alkyl chain attached at Q20 and Cy represents the alkyl chain attached N26. The x and y constitutes the number of carbons in the given alkyl chain. By this approach, we aimed at achieving a library of sCal peptides with a wide variety of lipid chain lengths allowing us to probe the mechanistic landscape of how lipidated sCal interacts with membranes and potentially bestow sCal with an intrinsic membrane translocating potential.

**Fig. 1.**
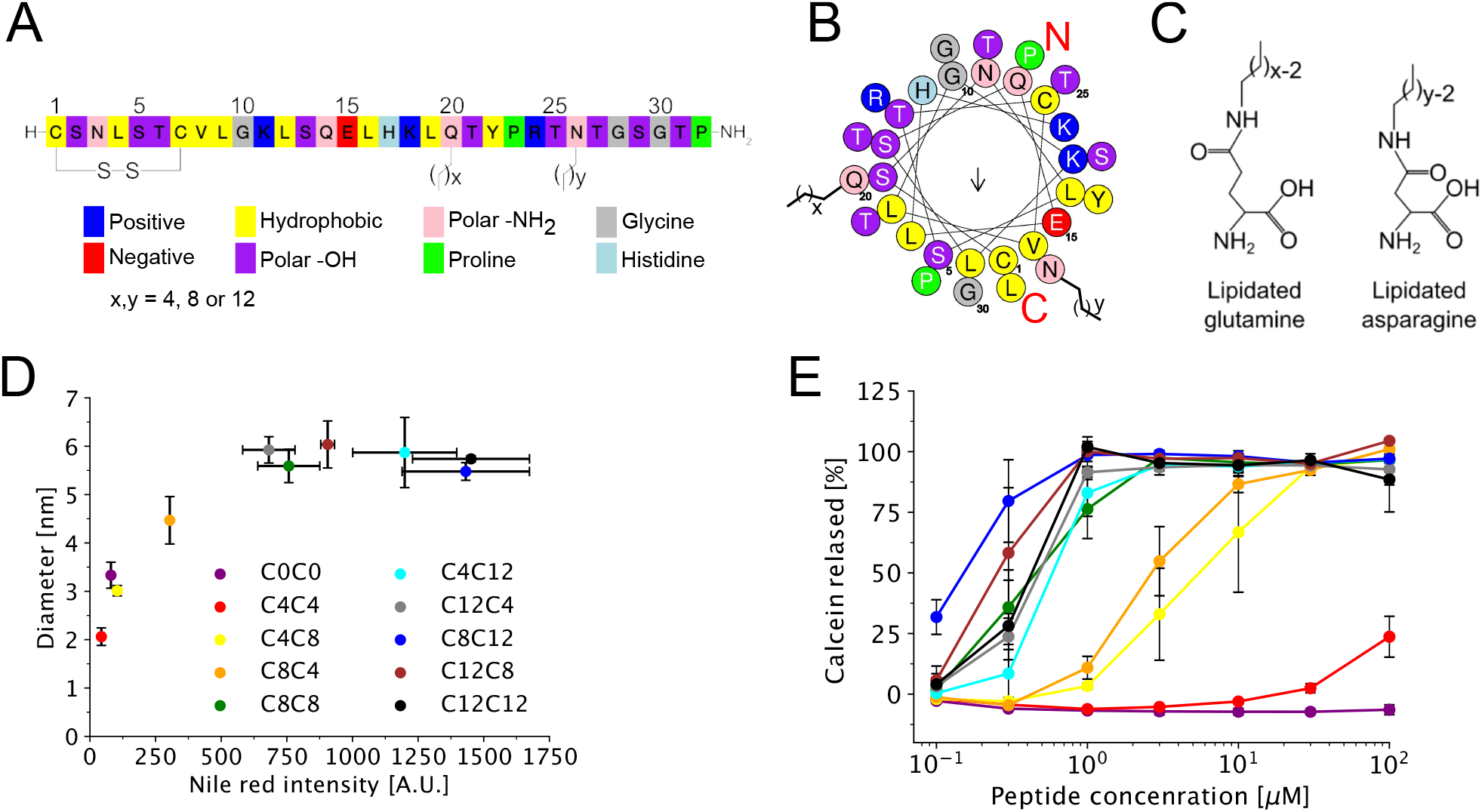
sCal peptide structure and bulk characterization of the average oligomer diameter, hydrophobicity and membrane activity of the lipidated sCal variants. A) Color-coded sCal amino acid sequence displaying cationic (dark blue), anionic (red), hydrophobic (yellow), polar (purple and pink), proline (green), glycine (gray) and histidine (light blue). Lipidated positions are at Q20 and N26 shown with stick representation. B) Helical wheel presentation of amino acid with N-terminal (red N), C-terminal (red C) and lipidation sites indicated (stick representation). The arrow inside the wheel represents the direction of the hydrophobic moment of native sCal (C0C0). The amino acid color scheme is the same as in A. C) Structures of synthesized lipidated glutamine and asparagine. D) The average diameter of the lipidated sCal variants, recorded by DLS, plotted against nile red intensity both measured at a peptide concentration of 100 μM. Data was collected from two independent experiments and error bars represent one standard deviation. All data points have y- and x-axis error bar, however some data points display error bars too small to be visible in the plot. E) Bulk calcein leakage from LUVs as a function of peptide concentration for lipidated sCal variants. The color scheme is the same as in D. Data was collected from two independent experiments and error bars represent the standard deviation. All data points have y-axis error bars, however some data points display error bars too small to be visible in the plot.

To characterize how lipidation affected the average oligomeric state of sCal we measured their size by DLS and hydrophobicity using a nile red assay (Fig. 1D). Overall, we found that native sCal (C0C0) and shorter lipidated sCal variants with up to a total of 12 carbons obtained an average diameter between 2 nm and 5 nm. Longer lipidated sCal variants, with a total of 16 carbons or more, obtained an oligomeric state with a stable average diameter of approximately 6 nm. The peptide with the smallest average diameter was not C0C0, but C4C4 with a diameter of 2.1 nm ± 0.2 nm. Assuming sCal is a spherical entity, the volume of a single non-lipidated peptide is 4.2 nm^3^, based on calculation from the structure, which in turn gives a diameter of approximately 2 nm (calculations found in supplementary information). This indicated that C4C4 was on average in a monomeric state, were as the other peptide variants, including C0C0, were on average in an oligomeric state. Interestingly, C8C4 and C4C8, representing analogs with identical total number of attached carbons, showed vastly different average diameters, indicating that the lipidation site influences peptide oligomerization (Fig. 1D). In the nile red assay we classified the sCal variants into four distinct hydrophobicity groups (Fig. 1D). See supplementary information for brief description of nile red assay. A low hydrophobicity group with C0C0, C4C4 and C4C8, a mid-low with C8C4, a mid-high with C8C8, C12C4 and C12C8 as well as a high with C4C12, C8C12 and C12C12. Considering the symmetrical sCal variants (C4C4, C8C8 and C12C12) we observed, as expected, an increase in hydrophobicity with increasing lipidation length. However, comparing the nile red intensity of an asymmetrical pair, e.g. C4C12 versus C12C4, vastly different intensities were recorded (Fig. 1D, cyan and grey). Together with the non-identical DLS diameters for other asymmetrical pairs, e.g. C8C4 versus C4C8, these data indicate that the hydrophobicity and oligomerization introduced by double lipidation cannot be considered only based on the total amount of lipid material added, but is represented by a complex interplay between lipidation site and length. These results highlight the importance of rigorous biophysical characterization of lipidated peptides to understand the structure-function relationships while studying membrane permeation.

### The membrane activity of sCal variants depends on lipidation length

To investigate how double lipidation affected sCal membrane activity, we measured calcein leakage experiments using LUVs encapsulating self-quenching concentrations of calcein and peptide concentrations ranging from 0.1 μM to 100 μM for each sCal variant (Fig. 1E). Overall, we found that sCal variants lipidated with 16 carbons or more in total were highly membrane active, all showing full calcein release in the low single digit μM range. sCal variants lipidated with 12 carbons in total were moderately membrane active displaying full calcein release above approx. 20 μM, while C4C4 displayed low activity not reaching full release for the tested peptide concentrations and C0C0 showed no membrane activity. Comparing C12C12 to C8C12 and C12C4 indicated that there was no increase in membrane activity when adding additional carbons beyond 16. This high-throughput quantitative analysis of membrane activity allowed us to sort the sCal variants into three distinct membrane activity groups, serving as the foundation for selecting a range of sCal variants for detailed mechanic studies.

### Time-lapse fluorescence microscopy of calcein flux into single GUVs reveals the membrane permeation mechanisms of sCal variants

The bulk calcein leakage assay provided an averaged membrane activity assessment, but did not reveal insights to the molecular mechanisms underlying the sCal mediated membrane permeation. Thus, we performed time-lapse recordings of GUVs in the presence of sCal variants as illustrated in Fig. 2A, using the influx of the fluorophore calcein into Atto655-DOPE labelled GUVs as a reporter for membrane activity. A time series of both the Atto655-DOPE and calcein channel were recorded for at least 30 minutes. A field of view (FOV) at t=0 is displayed in Fig. S2, illustrating how multiple GUVs were recorded per sample for each FOV. Each recorded GUV was tracked, quality assessed and analyzed using a custom python script (see methods and supplementary information for full description). In brief, 1) we corrected for an uneven illumination profile in the total FOV for each time frame, in each channel, 2) we localized each GUV using the membrane marker channel and 3) created a mask from the membrane marker channel allowing us to extract the mean intensity inside the GUV in the calcein channel for each frame, which 4) allowed us to calculate a ‘percent filled’ calcein value over time for each GUV, normalized to the mean local calcein background. The single GUV recordings and analysis provided an experimental framework for elucidating and comparing the kinetic membrane activity profiles for selected sCal variants.

**Fig. 2.**
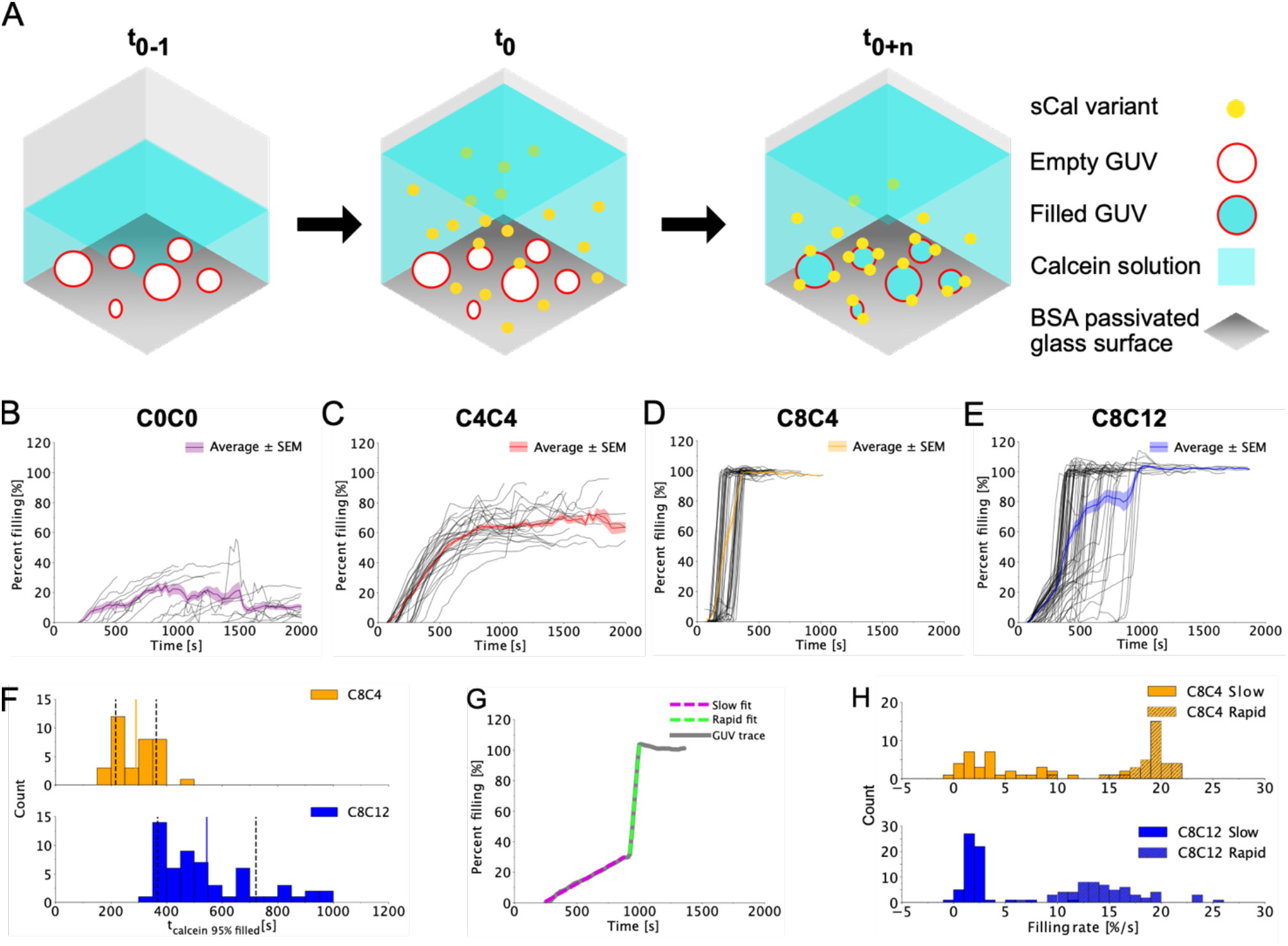
Single GUV time-lapse recordings of calcein influx reveal that variation of sCal lipidation strongly affects the kinetic membrane activity profile. A) Scheme depicting the setup of the single GUV experiments. GUVs are loaded into BSA coated wells, which have been filled to 50% capacity with a calcein solution at t_0-1_. Recording of the FOV are initiated at t_0_ after addition of a sCal peptide in a calcein solution, filling the well to 100% capacity and ensuring proper mixing. B-E) Percent calcein filling as a function of time for each sCal variant, C0C0 (B), C4C4 (C), C8C4 (D), C8C12 (E). The gray lines represent the quantification for each individual GUV trace, and the colored line represents the mean of all GUV traces. Note that the x-axis for B-E) range from 0-2000 s. The thickness of the colored line represents the standard error of the mean. Data was collected from two independent imaging sessions applying two individual GUV preparations with the following total number of wells: N_C0C0_ = 4, N_C4C4_ = 5, N_C8C4_ = 6 and N_C8C12_ = 5. F) Histogram of the time spent until a GUV is 95% filled with calcein for lapidated sCal variants C8C4 (yellow) and C8C12 (blue). Solid line displays average time spent for the GUVs reaching 95 % filled, while black dashed lines indicate ± one standard deviation. G) A representative trace for calcein filling percentage over time for a single GUV incubated with C8C12 (gray) fitted with a linear function for the two domains quantifying the slow filling rate (magenta) and the rapid filling rate (green). H) Histogram of filling rate values for slow (solid bars) and rapid (striped bars) rates for each GUV trace for lipidated sCal variants C8C4 (yellow) and C8C12 (blue).

### Small variations in sCal lipidation length strongly affect the kinetic profile of calcein filling into GUVs

We selected four lipidated sCal variants for analysis in the GUV assay, one from each of the membrane activity groups determined in the bulk calcein leakage data: A highly active (C8C12), a moderately active (C8C4), a slightly active (C4C4) and a non-active (C0C0) sCal variant (Fig. 1E). We compared the quantified calcein GUV lumen intensity values with that in the local background outside the GUV to calculate the percent calcein filling for each individual GUV (see method section for full description). We performed this calculation as a function of time for the selected sCal variants, with each gray line in figure 2B-E displaying the filling trace for one GUV. The individual filling traces for the same sCal variant were overall similar, producing a rolling average kinetic profile with a small standard error of the mean (SEM). On the contrary, when we compared the filling traces between the sCal variants we found distinct differences in the kinetic profiles. C0C0 and C4C4 displayed a single filling rate, both plateauing after 25 minutes at an average filling percent of approximately 20% and 65%, respectively (Fig. 2B and C). Control experiments with no peptides added show average GUV filling below 10% (Fig. S3). The C8C4 and C8C12 variants on the other hand display kinetic profiles with two filling rates, first a slow filling rate followed by a rapid filling rate (Fig. 2D and E). Visual inspection of traces reveals that the C8C12 traces in general maintained the slow filling rate for a longer period of time compared to C8C4, before switching to a rapid filling rate (see representative traces in Fig. S4). For both C8C4 and C8C12 the observed rapid rate ultimately lead GUV filling to 100% within our experimental time frame of 45 minutes. In conclusion, even minor alteration in lipid length e.g., from C4C4 to C8C4, leads to drastic difference in the kinetic calcein filling profile, suggesting that lipidation is a strong regulator of the intrinsic capacity of sCal to introduce membrane perturbations.

### Varying double lipidation characteristics strongly influences both slow rate duration and rapid rate kinetics

To quantify the filling kinetics for each sCal variant, we extracted the time between t_0_ and the time where the GUV was 95% filled (t_calcein 95% filled_) (Fig. 2F and Table 1). For the control and C0C0, no GUVs ever reached 95 % filling, while only 1 out of the 31 observed C4C4 GUVs reached 95% filled within the 45-minute timeframe of our experiment (Fig. S5). In contrast 100% of the C8C4 GUVs and 97 % of the C8C12 reached the 95% filled limit (Fig 2D-E and Table 1). Comparing the t_calcein 95% filled_ distributions for C8C4 and C8C12 revealed that C8C4 obtained a significantly higher t_calcein 95% filled_ distribution than C8C12 (CVM test, p-value < 0.001). The faster filling time for C8C4, compared to C8C12, remained at lower sCal peptide concentrations of 30 μM (Fig. S6). Next we wanted to understand which underlying features provided the basis for the faster C8C4 versus C8C12 GUV filling. This could be due to a shorter slow filling period or a higher rapid filling rate for C8C4. To quantify the time spent in the slow filling period, we plotted a distribution for the time from t_0_ till the tipping point of each GUV trace for C8C4 and C8C12 (Fig. S7). The average of each distribution displayed clearly that C8C4 spent significantly less time in a slow rate phase than C8C12 (CVM test, value < 0.001), with an average and one standard deviation values of 225 ± 68 s and 475 ± 167 s, respectively.

**Table 1.** Quantification of GUV filling, filling rates and observed GUVs for each sCal variant.

|  | Percentage of GUVs reaching 95 % filled - [%] | Average time before GUV reached 95 % filled – [s] | Slow rate – [%/s] | Rapid rate – [%/s] | Total number of GUVs |
| --- | --- | --- | --- | --- | --- |
| Control | 0 ± 0 | 0 ± 0 | 0.0 ± 0.3 | N.D. | 39 |
| C0C0 | 0 ± 0 | 0 ± 0 | 0.5 ± 0.4 | N.D. | 30 |
| C4C4 | 3 ± 4 | 1772 ± 0 | 2.1 ± 0.5 | N.D. | 31 |
| C8C4 | 100 ± 0 | 290 ± 73 | 4.6 ± 4.2 | 18.9 ± 2.2 | 35 |
| C8C12 | 97 ± 2 | 545 ± 177 | 2.2 ± 1.6 | 14.7 ± 3.7 | 59 |

Values are average ± one standard deviation. Control, C0C0 and C4C4 did not display rapid rate
behavior and therefore were not determined (N.D.).

To investigate the slow and rapid rates, we fitted a linear regression to the calcein trace for each GUV, determining percent filling per second. For C0C0 and C4C4 we fitted a single rate (Fig. S4), while for C8C4 and C8C12 we fitted individual linear regressions to both the slow and rapid rate (Fig. 2G). Examples of fitted rates are shown in Fig. S4. The mean ± SD of the slow distribution rates for the selected sCal variants were found to be 0.5 ± 0.4 %/s for C0C0, 2.1 ± 0.5 %/s for C4C4, 4.6 ± 4.2%/s for C8C4 and 2.2 ± 1.6 %/s for C8C12 (Fig. S8). CVM analysis showed that C0C0 obtained a significantly lower slow rate than the other sCal variants, while C8C4 obtained a significantly higher slow rate (Table S3). C4C4 and C8C12 were the only pair which were not significantly different (CVM test, p-value = 0.19) and obtained slow rates between C0C0 and C8C4. The significantly lower slow rate distribution for C0C0 compared to the lipidated variants suggests that lipidation overall increases the slow rate of sCal. For the rapid rate of C8C4 and C8C12 a CVM test showed that C8C4 possesses a significantly higher rapid rate distribution than C8C12 (Fig. 2H) (CVM test, p-value = 0.004). The observed variation of the rapid rate distributions for C8C4 and C8C12 were not linked to differences in GUV sizes, which range from 7.5 μm to 25 μm (Fig. S9). These analysis illustrates how is it possible to elucidate detailed information relating to the peptide:membrane interactions, using the single GUV assay. Furthermore these analysis allowed us to determine that the high filling kinetics observed for C8C4 originates from both the highest slow- and rapid filling rates and from spending the least amount of time in the slow rate phase. Overall these findings further support how minor alterations in lipidation characteristics can strongly influence the membrane perturbation capacity of sCal.

### Probing for physical dimensions of the membrane destabilization indicates that C8C4 and C8C12 induce major membrane destabilizations

To investigate the membrane destabilization mechanism leading to the slow and rapid filling kinetics for C8C4 and C8C12, we expanded the GUV assay to a dual reporter system by combining calcein (hydrodynamic range of 7 Å, [55]) with TRITC-Dextran-4kDa (Dex, hydrodynamic radius of 14 Å, [56]) (Fig. 3A). This dual reporter single GUV analysis allowed us to delineate the size dependencies of the membrane destabilization in real-time. A representative trace of a GUV filling with C8C12 present in the dual reporter system is displayed in Fig. 3B (Fig. S10 for all traces). The calcein filling kinetics in the dual reporter system displayed similar profile characteristics as the calcein in the single reporter system, demonstrating that no large-scale interference occurred between calcein and Dex (compare Fig. 2B-E and S10). The Dex traces overall displayed a lag phase prior to transitioning directly to a rapid filling phase (Fig. 3B, orange). To quantify and compare the filling kinetics between calcein and Dex observed for C8C4 and C8C12, we extracted the time spent for a single GUV before reaching 95 % filled for both Dex (t_Dex 95% filled_) and calcein (t_calcein 95% filled_). For each GUV we calculated the temporal difference between t_calcein 95% filled_ and t_Dex 95% filled_ as Δt_95% filled_ = t_calcein 95% filled_ – t_Dex 95% filled_ and plotted these in a histogram (Fig. 3C). Positive values correspond to calcein reaching 95 % filled before Dex, while negative represent the opposite. For both C8C4 and C8C12 only positive values are observed, meaning calcein always filled the GUV to 95 % before Dex. This temporal difference could be caused by a difference in diffusion coefficient between calcein and Dex. Using the Stokes-Einstein equation for diffusion of a spherical particle in a liquid, the diffusion coefficients of calcein and Dex were determined in water to be 326.0 μm^2^/s and 185.6 μm^2^/s, respectively. Calcein hereby diffuses 57% faster than Dex, however when considering that the GUV sizes range between approximately 7.5 μm to 25 μm, the difference in diffusion is negligible within our temporal imaging resolution of 15 s. The temporal difference between the two reporters for both C8C4 and C8C12 indicated that the peptide:membrane interactions initially induce minor perturbations of the bilayer, only allowing transport of the smaller calcein across the membrane occurring during the slow rate phase. Subsequently, as the calcein filling profile transitions to the rapid rate, this also leads to the influx of Dex, suggesting a major membrane destabilization consistent with the formation of membrane pores.

**Fig. 3.**
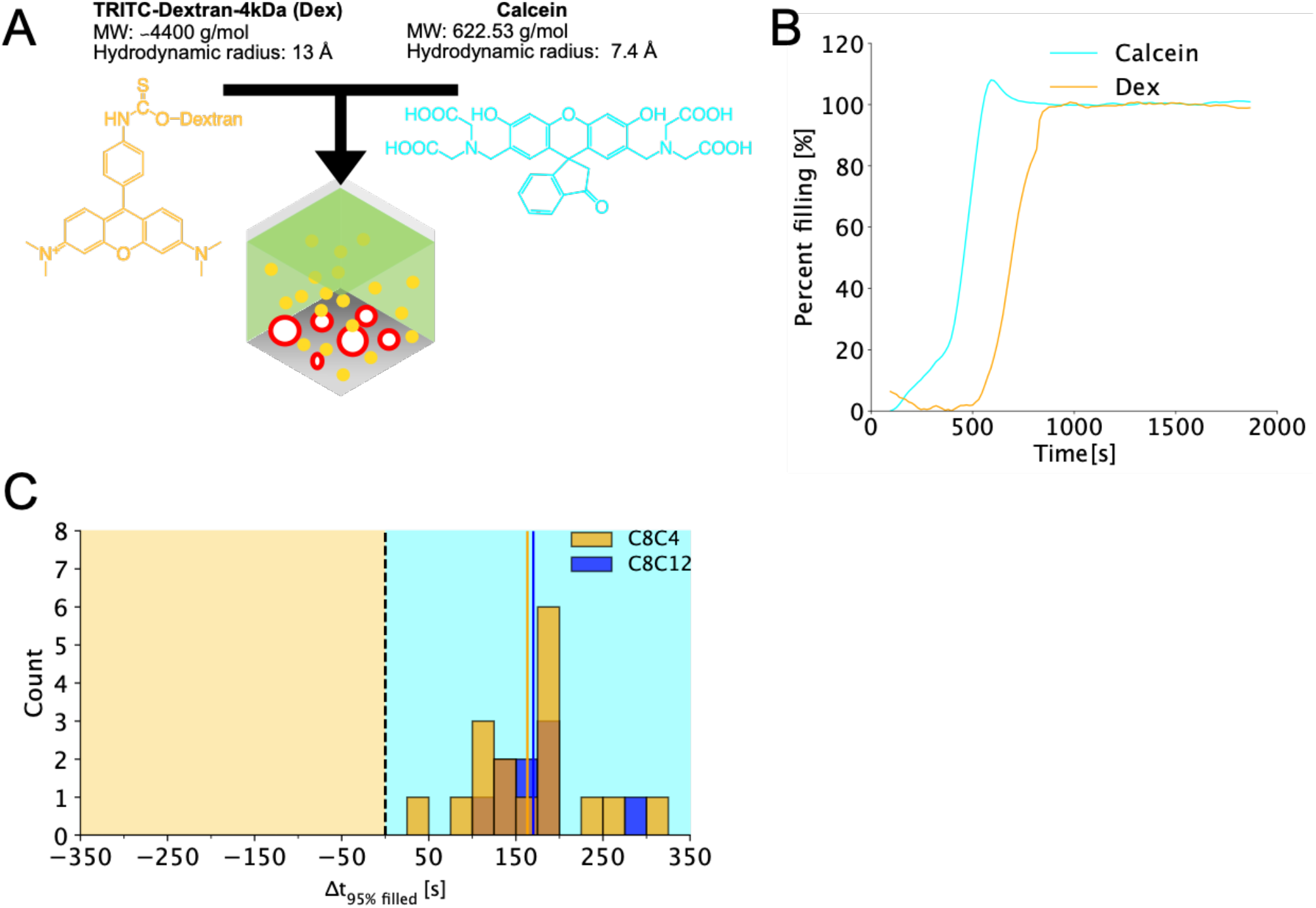
Dual reporter system probing the size characteristics of peptide mediated membrane instabilities suggests that lipidated sCal can form pores. A) Illustration of the GUV setup including two reporter molecules of different size and emission profiles: Calcein and Dex. B) A representative trace for calcein (cyan) and Dex (orange) depicting the filling percentage over time for a single GUV incubated with C8C12. C) Histogram showing the temporal difference between the time from when, for the same GUV, calcein reaches 95% filled to when Dex reaches 95% filled, when incubated with C8C4 (yellow) or C8C12 (blue). Data was collected from two independent imaging sessions applying two individual GUV preparations with the following amount of wells: N_C8C4_ = 8 and N_C8C12_ = 6.

### GiGs assay suggests that sCal membrane translocation requires a peptide mediated pore forming mechanism

Until now, we have focused on the membrane perturbation mechanism induced by lipidating sCal. Next we wanted to investigate how these properties were linked with peptide membrane translocation. To do this we employed a GiGs based assay utilizing the time-lapse microscopy recordings of calcein influx (Fig. 4A). By monitoring the filling of both the outer and the inner GUV we were able to distinguish peptide mediated membrane perturbation events that only leads to calcein influx into the outer GUV from events where both the outer and inner GUV are filled. Since the inner GUV will only show calcein influx if the peptides are able to translocate across the membrane of the outer GUV, hereby reaching the membrane of the inner GUV, we can use this setup to determine the propensity of lipidated sCal variants to translocate across membranes. Importantly, this assay allows for studying peptide translocation without the need for fluorescently labelling the peptide, which has been shown to alter the physicochemical properties of peptides and hereby their intrinsic membrane interaction characteristics [44]. We manually localized suitable GiGs in the time-lapse recordings and monitored the calcein filling of both the inner and outer GUV (Fig. 4B-E). Distinct behaviors for the four sCal variants were evident as C0C0 showed no GiGs with even a significant filling of the outer GUV, consistent with the data recorded for the single GUV setup. For C4C4 the prevailing behavior consisted of GiGs displaying a significant influx of calcein in the outer GUV, but very limited influx in the inner GUV. Finally, both C8C4 and C8C12 displayed strong calcein filling of both the outer and inner GUV. To quantify the behavior of the sCal variants in the GiG setup we monitored at least 50 GiGs for each peptide variant and calculated the percentage of GiG per chamber that displayed at least 60% filling of the outer- and inner GUV (Fig. 4F and Table S4). We found that neither the control nor the C0C0 data contain any GiG displaying significant filling in either the inner or the outer GUV. For the C4C4 peptide we quantified that 54 ± 8% of the outer GUVs filled, but only 16 ± 13% displayed filling of the inner GUV. On the contrary, all GiGs monitored for the C8C4 and C8C12 displayed filling of both outer- and inner GUV. This data demonstrates that only C8C4 and C8C12 showed substantial peptide translocation across membranes. This correlates with these two peptides both exhibiting rapid rate kinetic profiles, suggesting that the large scale pore-like perturbation associated with this phase is essential for peptide translocation.

**Fig. 4.**
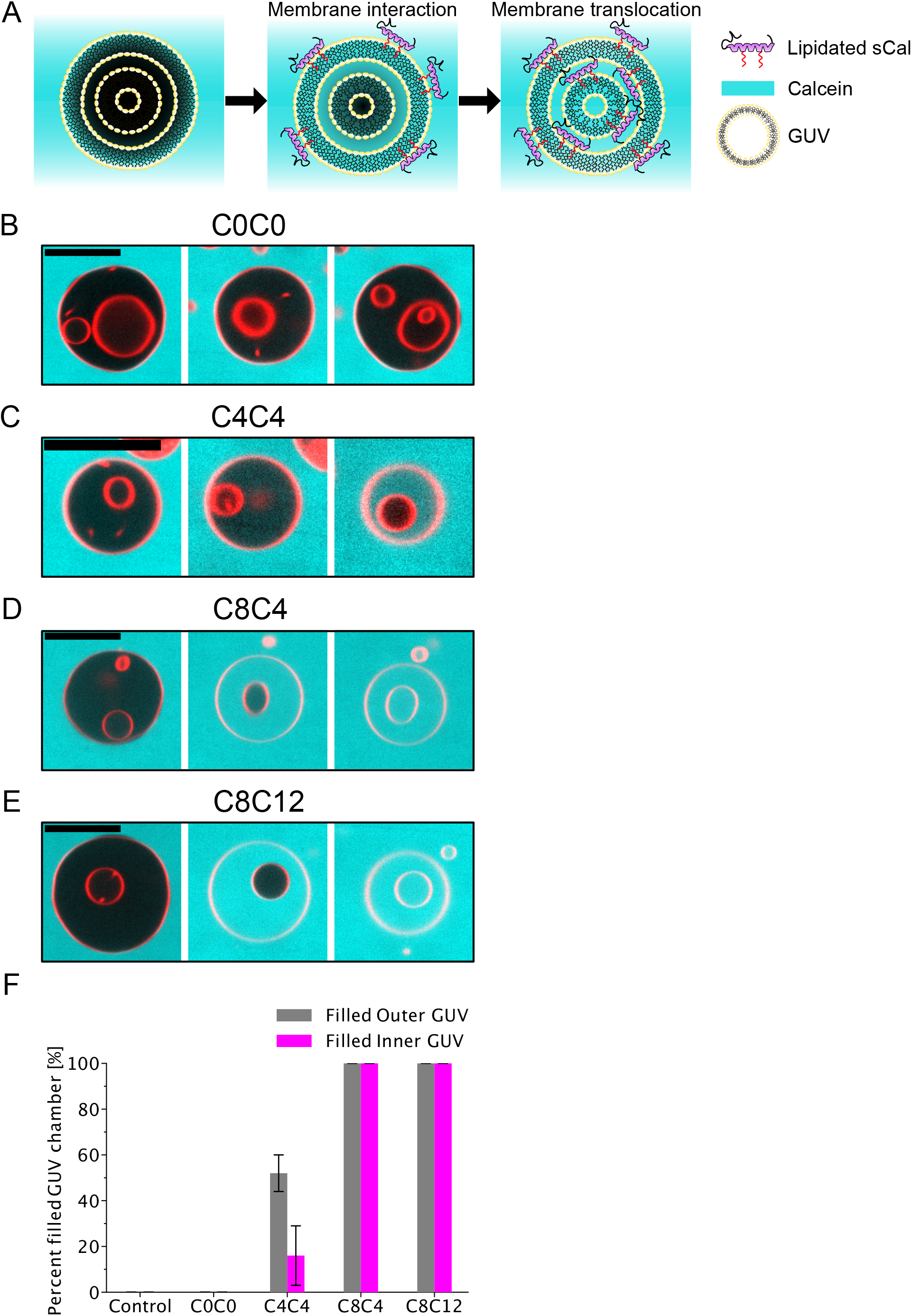
Monitoring calcein influx in GiGs reveals a lipidation-dependent propensity of sCal to translocate across membranes. A) Scheme depicting the possible outcomes of the GiG experiments. A sCal variant only displays membrane translocation properties if both the outer- and the inner GUV is filled with calcein. B-E) Micrographs from three different time points for the same GiG showing the GUV membrane (red) and calcein (cyan) signals for C0C0 (B), C4C4 (C), C8C4 (D) or C8C12 (E). Scale bar is 20 μm in all micrographs. F) Bar plots showing the percent of GiGs per chamber displaying calcein filling of the outer (gray) and inner (magenta) GUVs Data was collected from two independent imaging sessions applying two individual GUV preparations with the following amount of wells: N_C0C0_ = 4, N_C4C4_ = 5, N_C8C4_ = 6 and N_C8C12_ = 5. Error bars represent one standard error of the mean.

## Discussion

The wide range of employed assays and results presented in this study, from determining bulk biophysical peptide properties to imaging of membrane perturbation mechanisms, provided a robust peptide characterization, which importantly did not require peptide labeling. In combination, the assays created a broadly applicable workflow, delivering valuable insights to the membrane interaction mechanism of peptides. First, we employed rigorous biophysical characterization techniques to investigate how double lipidation of the model therapeutic peptide, sCal, affected properties like oligomeric state and membrane activity. Understanding and controlling the oligomerization of lipidated peptides are paramount for their successful application in oral delivery, as underscored by the first clinically approved oral formulation for treatment of diabetes Rybelsus [2,57,58]. This formulation consists of a lipidated peptide, semaglutide, co-formulated with the permeation enhancer salcaprozate sodium (SNAC). One of the main roles of SNAC is to ensure semaglutide is in a monomeric form, which facilitates its transport across the cellular barrier [59].

We observed that for sCal peptides with longer chains, from 16-24 total carbons, a maximum size of approximately 6 nm was reached, indicating that an upper limit of peptide oligomerization. Comparing the short chain sCal peptide C4C4, to native sCal, C0C0, we saw that while C4C4 displayed a size matching the monomer, the C0C0 appeared to form an oligomeric structure. This ability of lipidation to induce a more monomeric state is corroborated by the work of Wang *et al.* [60] which measured the size of native and single palmitic acid lipidated glucagon-like peptide-1/glucose-dependent insulinotropic polypeptide hybrid peptides. Depending on peptide concentration, they observed smaller sizes for the lipidated peptide compared to the native peptide. Further, the observation that C4C8 and C8C4 display distinct sizes, indicating that not only the amount, but also the placement within the peptide structure can greatly influence oligomerization propensity of peptides, supports that lipidation of peptides can lead to hard-to-predict structural effects. Taken together our data support that intrinsically assuming that peptide oligomerization simply scales with the amount of hydrophobicity introduced by e.g., lipidation can be faulty and thus the oligomerization state for modified peptides should always be assessed experimentally.

Comparing the membrane activity of the symmetrical sCal variants, C0C0, C4C4, C8C8 and C12C12 in our calcein leakage study (Fig. 1E) we found a proportionality between increasing amount of lipidation and increasing membrane activity. However, by including the asymmetrical sCal variants we discovered that the most potent sCal variant was C8C12, which at the lowest measured peptide concentration (0.1 μM) obtained an approximately 7 folder higher membrane activity than the sCal variant with the most added carbons, C12C12. C8C12 and C12C12 obtained similar average oligomeric sizes and hydrophobicities, indicating that that the membrane activity bestowed by lipidation cannot easily be predicted from classical biophysical characterization parameters.

Next, we used single GUV measurements to extract how lipidation alters the molecular mechanisms of sCal mediated membrane destabilization and further how this relates to peptide translocation. Previous efforts using single GUV calcein influx setups performing endpoint measurement on membrane-active peptides have allowed researchers to assign such peptides to harbor either an AoN and graded behavior [23,27]. Relating this to our data we saw that at the end of the measurement (after 45 minutes) we have sCal variants displaying only a graded mechanism (C0C0 and C4C4), while others show both a graded and an AoN mechanism (C8C4 and C8C12). This suggests that even minor changes in the lipid chain lengths can alter the peptide mediated perturbation mechanism. The AoN perturbation mechanism is, as previously described, associated with pore formation, while graded filling is defined to originate from numerous weak and short-lived membrane perturbations [37]. This is interesting when we investigate the time-dependent change in the filling rates seen for the two most potent sCal variants C8C4 and C8C12. We found that C8C12 obtains comparable slow rate kinetics to C4C4 (Fig. S8), while C8C4 obtains a slightly higher slow rate. Even though C8C4 has a marginally higher slow rate, these results suggest that initially the GUVs are experiencing graded filling, however upon switching to the rapid rate, an AoN filling mechanism is experienced. Such a lipidation dependent switch in the membrane destabilization mechanism over time could potentially be linked with the build-up of sCal on the surface of the GUV reaching a threshold value that switches the filling behavior from graded to AoN. A dual behavior in GUV filling kinetics has previously been seen for magainin 2, a membrane-active peptide, which showed an initial minor release of calcein from GUVs followed by a rapid leakage [40]. The rapid leakage for magainin 2 was attributed to the onset of pore formation, which further supports that the rapid filling phase seen in our GUV assay could also originate from the formation of sCal-mediated membrane pores. We also note that macrolittin-70, a well described pore forming membrane active peptide [61], tested in our single GUV assay displayed a rapid rate phase calcein filling profile very similar as for the C8C4 and C8C12 sCal variants (Fig. S11). Furthermore, we observed that the rapid phase filling of GUVs happens relatively fast, sometimes within 1.5 minutes, nonetheless GUVs stayed intact and could be tracked until complete filling has occurred (Fig. S4). This again suggests a mechanism not based on micellization of the lipid membrane, but a pore forming scenario where the GUV bilayer maintained its overall integrity. The data produced for the dual reporter system further supports the radical switch in filling behavior seen for the C8C4 and C8C12 sCal variants. Combining the visual evaluation of the calcein and Dex traces with the quantitative assessment of the filling indicated that the peptide:membrane interactions initially induce minor perturbations of the bilayer, only allowing transport of the smaller calcein across the membrane. As the calcein filling profile transitioned to the rapid phase, this also led to the influx of the larger Dex molecules, suggesting a shift in the size of the membrane destabilizations compatible with the notion that going from slow to rapid phase constitutes a change to a pore-mediated filling mechanism. Together the data shown support that fine-tuning the double lipidation of sCal can bestow the peptide with an intrinsic pore-forming ability that could greatly enhance the cross barrier delivery.

## Conclusions

In this study, we used time-lapse fluorescent imaging of single GUVs to elucidate the mechanistic details of how lipidating therapeutic peptide affected their membrane perturbation mechanism. We found that even minor modifications in lipid chain lengths can alter peptide mediated membrane destabilizations from being based on many weak and short-lived membrane perturbations to being based on peptide-pores. Introducing the most potent pore formation did not simply correlate with increasing lipid chain length, underscoring the complex interplay between oligomerization and membrane interaction established when lipidating peptides. Interestingly, only the sCal variants which obtained pore forming properties displayed efficient membrane translocation. The results presented here illustrate how investigating the molecular mechanism of peptide function can reveal novel peptide:membrane interaction characteristics that could serve as the foundation for further peptide reengineering. Our findings for sCal presented double lipidation as a valuable strategy as an intrinsic peptide permeation enhancer to facilitate the cross barrier transport of orally delivered therapeutic peptides.

## Author contributions

P.M.L. and J.B.L. designed research; P.M.L., K.K., N.W.L. and A.Z.E. performed experiments; A.K., M.B.H. and C.U.H. performed peptide synthesis and purification; P.M.L created analytic tools with assistance from K.M.; P.M.L. performed data analysis; A.J.U., J.B.S., J.B.L. and T.L.A. supervised project; P.M.L. and J.B.L. wrote the paper. All authors discussed the results and commented on the manuscript.

## Acknowledgements

The authors thank Fredrik Melander for performing the ICP-MS recordings. This work was supported by the Novo Nordisk Foundation (grant no. NNF16OC0022166 to TLA). The data that support the findings of this study and the custom python scripts are available from the corresponding author upon reasonable request.

