## Supplementary information for "Tuning the double lipidation of salmon calcitonin to introduce a pore-like membrane translocation mechanism"

#### Experimentals for amide-functionalized sCal library production

All standard and orthogonally protected $N_{\alpha}$-Fmoc amino acids were purchased from Iris-Biotech. $Pd(PPh_3)_4$ was purchased from Alfa Aesar, and all other reagents were purchased from Sigma Aldrich. NMR Spectra were recorded on a 400MHz Bruker NMR spectrometer.

Peptides were synthesized on a Biotage Initiator + Alstra microwave assisted peptide synthesizer. Peptides were purified on a reverse-phase Waters HPLC system equipped with a 600 controller, 600 pumps and a UV/Vis 2489 detector (monitoring at 214nm and 280nm) using a Waters Xbridge C18 OBD, 5 um, 250 x 19 mm column. Gradients were run using a solvent system consisting of A (5% MeCN in $H_2O$ + 0.1 % TFA) and B (MeCN + 0.1 % TFA), and collected fractions were lyophilized on a Labogene Scanvac Coolsafe (-110 $^{o}$C).

Pure peptides were analyzed on a Shimadzu NexeraX2 reverse-phase HPLC (RP-HPLC) system equipped with Shimadzu LC-30AD pumps, a Shimadzu SIL-30AC autosampler, a CTO-20AC column oven and a Shimadzu PDA detector (monitoring at 214nm and 280nm) using a Waters XBridge BEH C18, 2.5µm 3.0x150mm XP Column at a flow rate of 0.5 mL/min. RP-HPLC gradients were run using a solvent system consisting of solution A (5% MeCN in $H_2O$ + 0.1% TFA) and B (MeCN + 0.1% TFA). Analytical HPLC was using to characterize each peptide; a gradient from 25% to 75% solution B over 25 min. Analytical RP-HPLC data is reported as column retention time ($t_R$) in minutes (min) (Fig S12-S21). Low resolution mass spectrometry (LRMS) was performed on a Bruker MALDI-TOF Autoflex speed using Flexcontrol 3.4 software.

Peptide content was analyzed on a Nanodrop 2000c using UV absorption of peptides at 280nm (extinction coefficient 1615 AU/mmol/ml)

#### Peptide synthesis

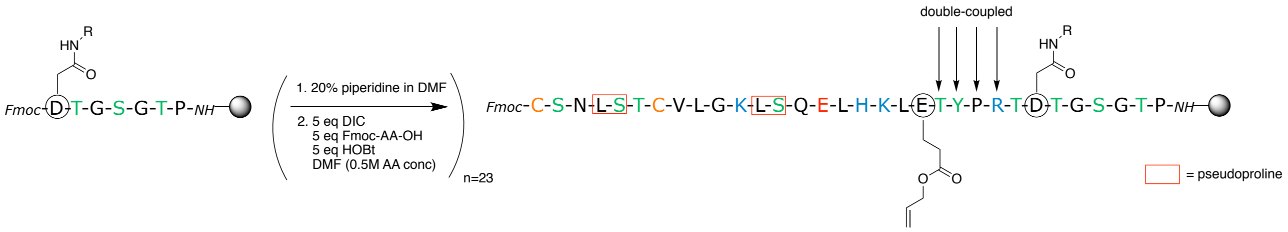

Peptides were synthesized using Tentagel S RAM amide Resin (0.24 mmol/g) (Rapp-Polymer). Couplings were performed using either 4 equivalents Fmoc-protected amino acid/4 equivalents HCTU/4 equivalents DIPEA in DMF (3 mL), or 5 equivalents Fmoc-protected amino acid/ 5 equivalents DIC/5 equivalents HOBt. Coupling of standard Fmoc-protected amino acids was carried out for 5 min at 75 $^{o}$C followed by 4x 45s washes. Arginine was double coupled; 60 min at room temperature followed by 5 min at 75 $^{o}$C, and repeated with fresh reagents followed by washing. Histidine and cysteine were coupled at 50 $^{o}$C for 10 min followed by washes. Deprotection was carried out in 20% piperidine in DMF + 5% formic acid for 30s and then 3 min at room temperature followed by washing.

Special synthesized amino acids ($N_4$, $N_8$ or $N_{12}$; see synthesis below) were incorporated using standard synthesis on the Alstra (**D** in sequence above), followed by incorporation of Fmoc-L-Glu(OAll)-OH. After final automated synthesis, the batch was split in three and for each batch the OAll was removed and the respective amine chains were coupled as specified below.

Test cleavages were performed in a cleavage cocktail (1 mL) of 94% TFA, 2.5% $H_2O$, 2.5% 1,2-ethanedithiol and 1% triisopropylsilane for 30 min at room temperature and the cleavage cocktail was evaporated using a stream of nitrogen. The peptide was precipitated from solution with ice cold $Et_2O$, centrifuged at 4500 rpm for 5 min and the precipitate was dissolved in 50/50 $MeCN/H_2O$, centrifuged at 4500 rpm for 5 min and run using MALDI-TOF and analytical HPLC.

Peptides were cleaved in a cleavage cocktail (10 mL) of 94% TFA, 2.5% EDT, 2.5% $H_2O$ and 1% triisopropylsilane. The resin was stirred for 2.5 hours at room temperature and the cleavage cocktail was evaporated using a stream of nitrogen. The peptides were precipitated from solution with ice cold $Et_2O$, centrifuged at 4500 rpm for 5 min and the precipitate washed with ice cold $Et_2O$. The peptide was dissolved in $H_2O$/MeCN + a few drops of AcOH and lyophilized overnight.

General method for OAll-removal

The peptide resin was swelled in dry $CH_2Cl_2$ and washed 2x in dry $CH_2Cl_2$.

The resin was then added 3 mL dry $CH_2Cl_2$ (slurry) and 24 eq phenylsilane and shaken for 5 min at room temperature. 0.25 eq $Pd(PPh_3)_4$ was then added in 200uL dry $CH_2Cl_2$. The resin was wrapped to extrude light and shaken carefully at room temperature for 40 min. The resin was then washed with dry $CH_2Cl_2$ and the full procedure was repeated.

The resin was washed in 3x $CH_2Cl_2$, 3x IPA, 3x DMF, 3x 0.5% sodium diethyldithiocarbamate in DMF, 3x $CH_2Cl_2$ and test-cleaved.

General method for coupling after OAll removal

The resin was swelled in DMF and coupled in 4 eq HATU/4 eq DIPEA/4 eq of the relevant amine carbon chain (butylamine/octylamine/dodecaneamine) for 1 hour at room temperature. The coupling was repeated with fresh reagents. The resin was then washed 3x DMF, 3x IPA, 3x $CH_2Cl_2$ and test-cleaved.

Iodine oxidation

Following full cleavage and lyophilization, the peptide was dissolved in a round-bottomed flask in 50/50 MeOH/$H_2O$ at a concentration of 1mg/mL (Note: the more hydrophobic peptides were dissolved in 2mL MeOH to start with). A stock solution of 0.06M $I_2$ in MeOH was added to the stirring peptide until a yellow color persisted. The excess iodine was then quenched by addition of a 1M ascorbic acid solution (in $H_2O$) until the solution was colorless. The mixture was then freeze-dried.

#### Amino acid synthesis

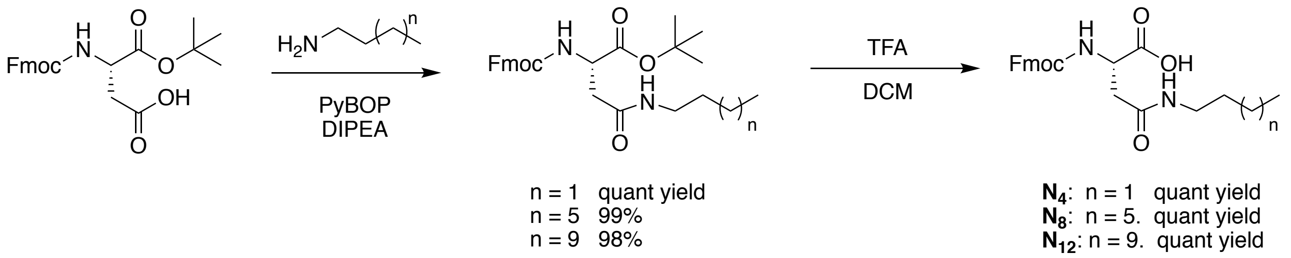

Coupling of fatty acid to Fmoc-Asp-OtBu

To a round-bottomed flask Fmoc-Asp-OtBu (316 mg, 0.77 mmol, 1.0 eq) was added and dissolved in $CH_2Cl_2$ (50 mL), followed by the addition of PyBOP (400 mg, 0.77 mmol, 1.0 eq), DIPEA (294 $\mu$L, 1.69 mmol, 2.2 eq) and finally butylamine/octylamine/dodecylamine (0.85 mmol, 1.1 eq). The solution was stirred at rt overnight. The $CH_2Cl_2$ was evaporated and the residue dissolved in EtOAc. The organic phase was washed with 2x50mL 1M $KHSO_4$, 2x50mL $NaHCO_3$(sat) and 2x50mL brine, dried with $MgSO_4$, filtered and concentrated *in vacuo*.

Ester-deprotection to make to make $N_4$, $N_8$ and $N_{12}$

The fatty acid amino acids were each dissolved in 10 mL $CH_2Cl_2$ under vigorous stirring and 5 mL TFA was added, and the reaction was stirred for 40 min at rt.

The solvent was then evaporated using a steady stream of $N_2$ gas, after which 50mL $CH_2Cl_2$ was added and the residue concentrated *in vacuo*, and the procedure was repeated twice more and the Fmoc-amino acids were isolated as a white powder.

#### Peptide characterization

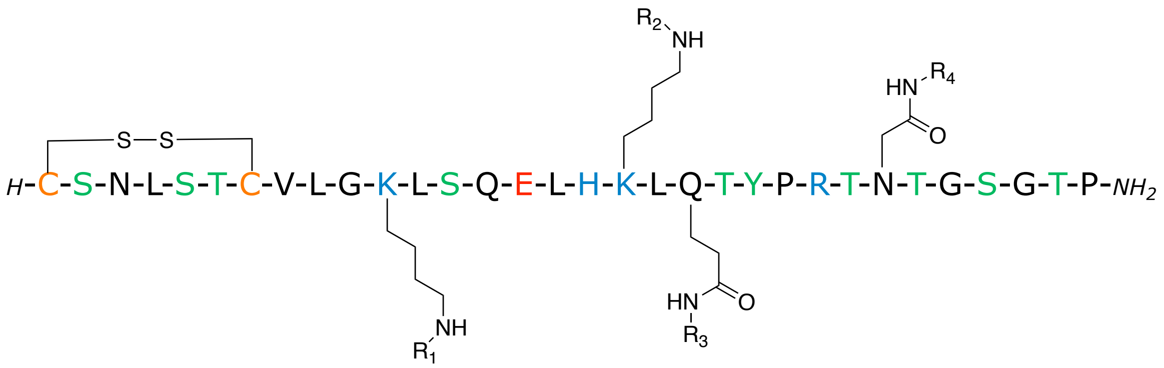

| Peptide | Structure | | | | $t_R$ (min) | Purity (214nm) | Yield |
|---|---|---|---|---|---|---|---|
| | $R_1$ | $R_2$ | $R_3$ | $R_4$ | | | |
| $C_0C_0$ | H | H | H | H | 8.90 | >99% | 16.1 mg (26%) |
| $C_4C_{12}$ | H | H | $-(CH_2)_3CH_3$ | $-(CH_2)_{11}CH_3$ | 17.20 | >99% | 19.7mg (24%) |
| $C_8C_{12}$ | H | H | $-(CH_2)_7CH_3$ | $-(CH_2)_{11}CH_3$ | 20.37 | >99% | 15.6mg (19%) |
| $C_{12}C_{12}$ | H | H | $-(CH_2)_{11}CH_3$ | $-(CH_2)_{11}CH_3$ | 24.19 | >99% | 18.1mg (22%) |
| $C_4C_4$ | H | H | $-(CH_2)_3CH_3$ | $-(CH_2)_3CH_3$ | 11.90 | >98% | 10.6mg (13%) |
| $C_8C_4$ | H | H | $-(CH_2)_7CH_3$ | $-(CH_2)_3CH_3$ | 14.92 | >99% | 18.4mg (23%) |
| $C_{12}C_4$ | H | H | $-(CH_2)_{11}CH_3$ | $-(CH_2)_3CH_3$ | 18.63 | >99% | 14.3mg (18%) |
| $C_4C_8$ | H | H | $-(CH_2)_3CH_3$ | $-(CH_2)_7CH_3$ | 13.98 | >98% | 13.0mg (16%) |
| $C_8C_8$ | H | H | $-(CH_2)_7CH_3$ | $-(CH_2)_7CH_3$ | 17.02 | >99% | 13.2mg (16%) |
| $C_{12}C_8$ | H | H | $-(CH_2)_{11}CH_3$ | $-(CH_2)_7CH_3$ | 20.60 | >98% | 5.6mg (7%) |

#### HPLC characterization

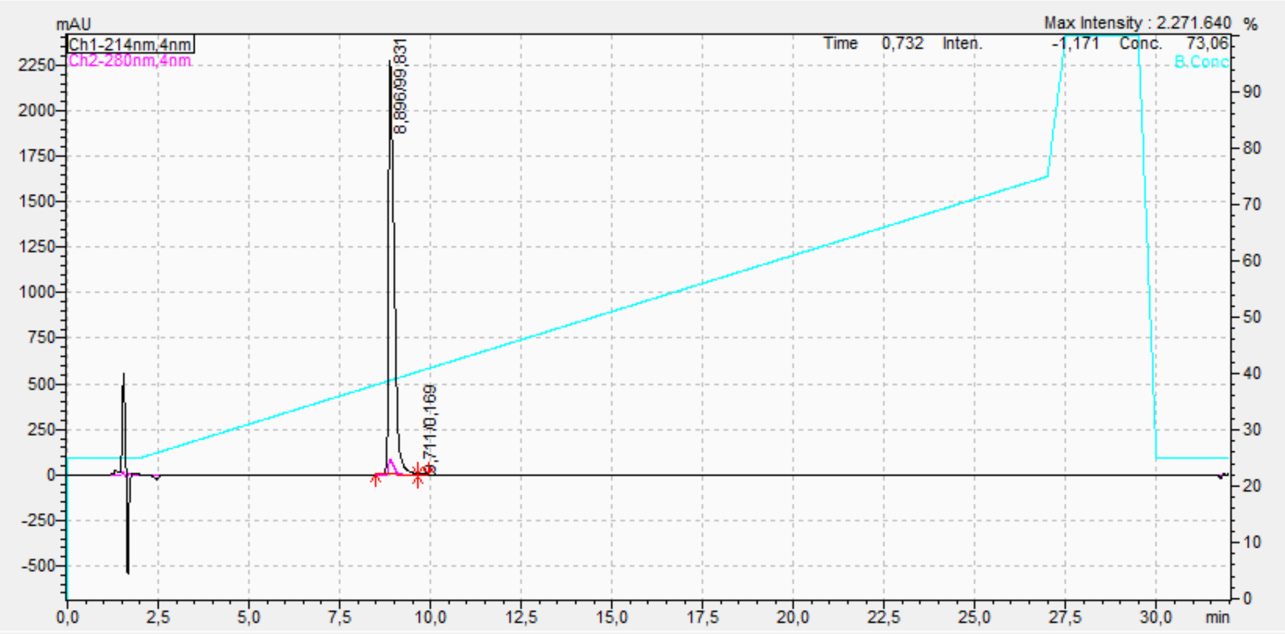

**C0C0 purity.** Analytical HPLC chromatogram.

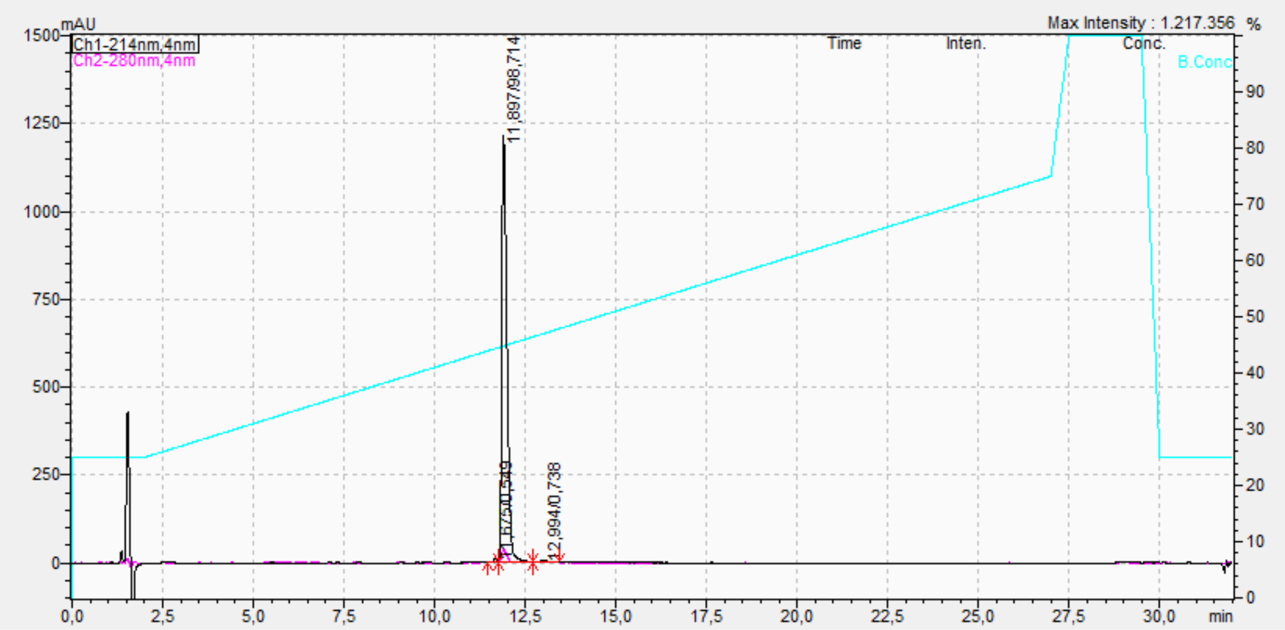

**C4C4 purity.** Analytical HPLC chromatogram.

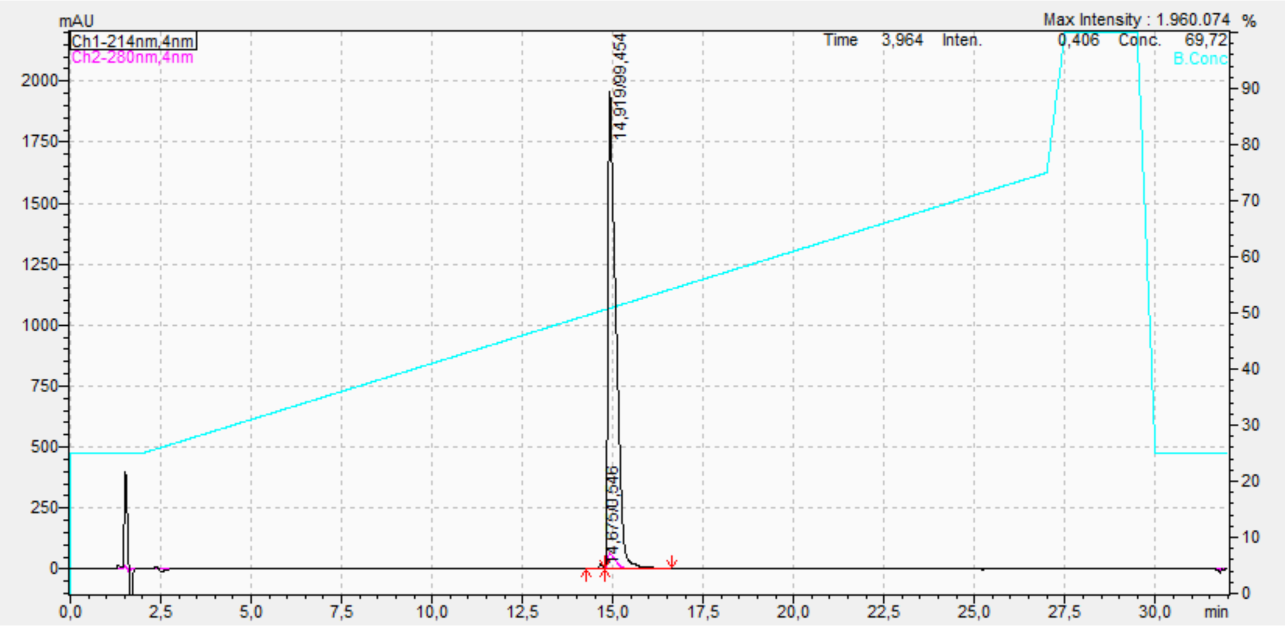

**C8C4 purity.** Analytical HPLC chromatogram.

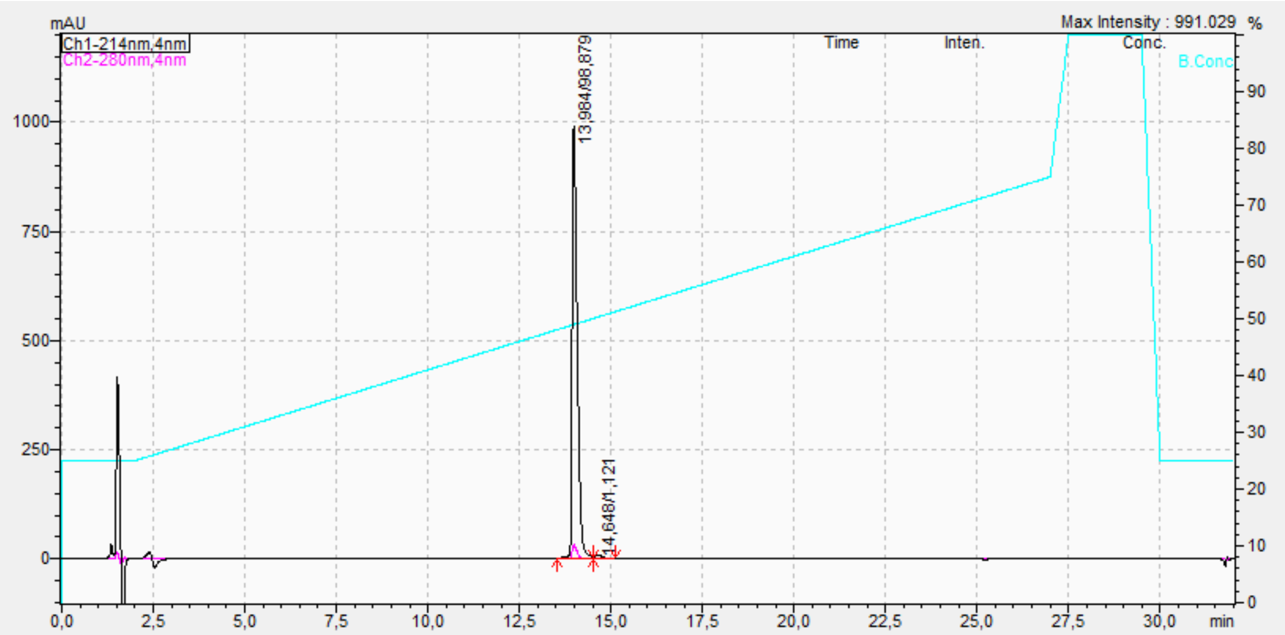

**C4C8 purity.** Analytical HPLC chromatogram.

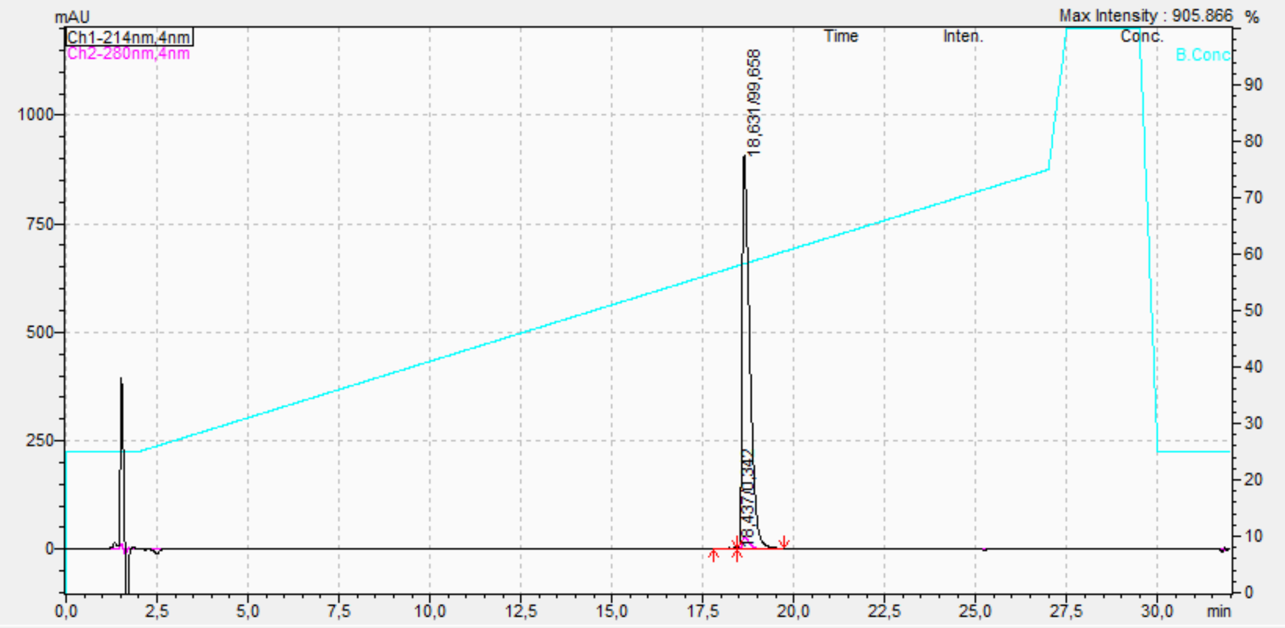

**C12C4 purity.** Analytical HPLC chromatogram.

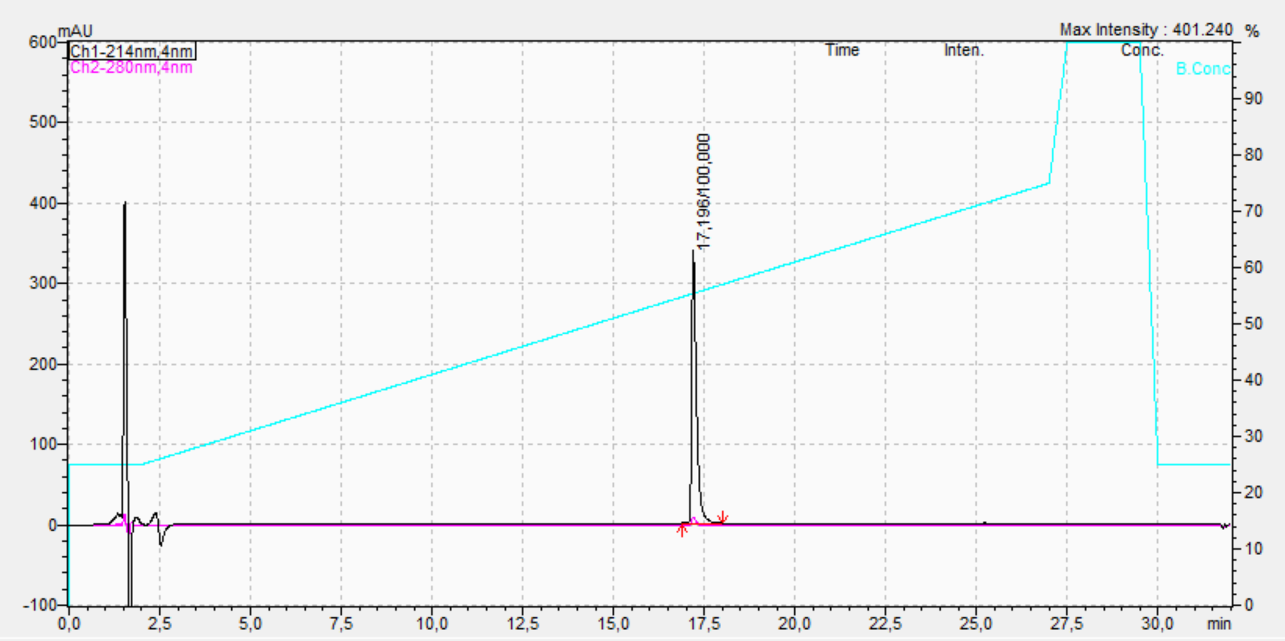

**C4C12 purity.** Analytical HPLC chromatogram.

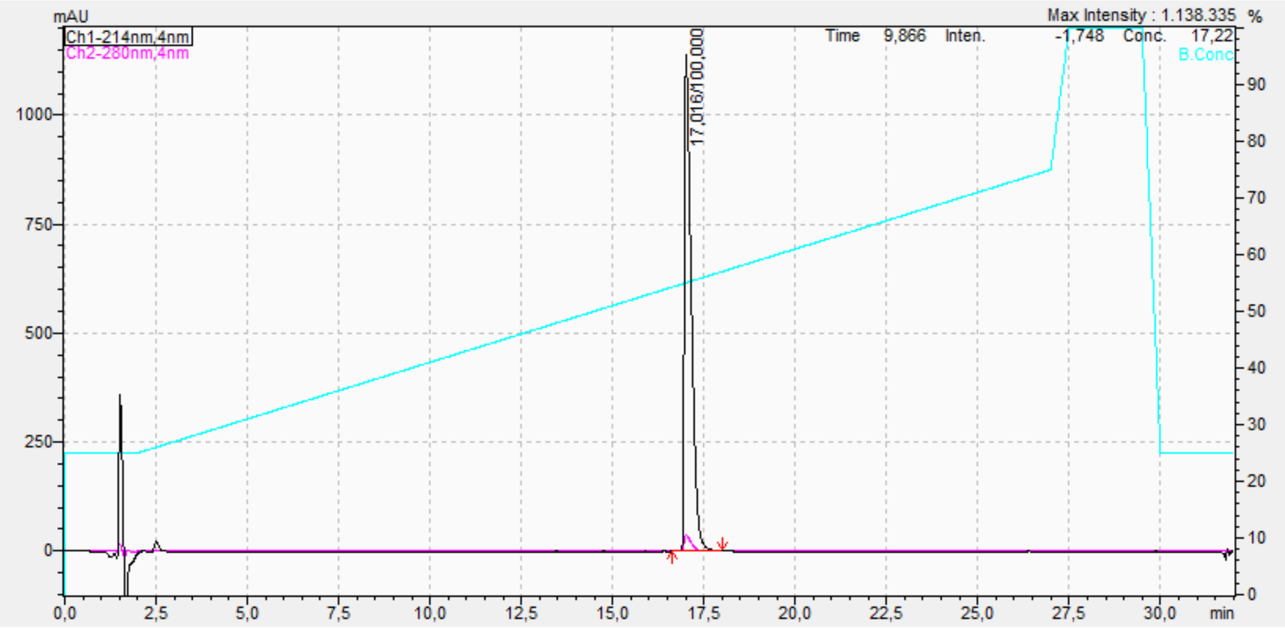

**C8C8 purity.** Analytical HPLC chromatogram.

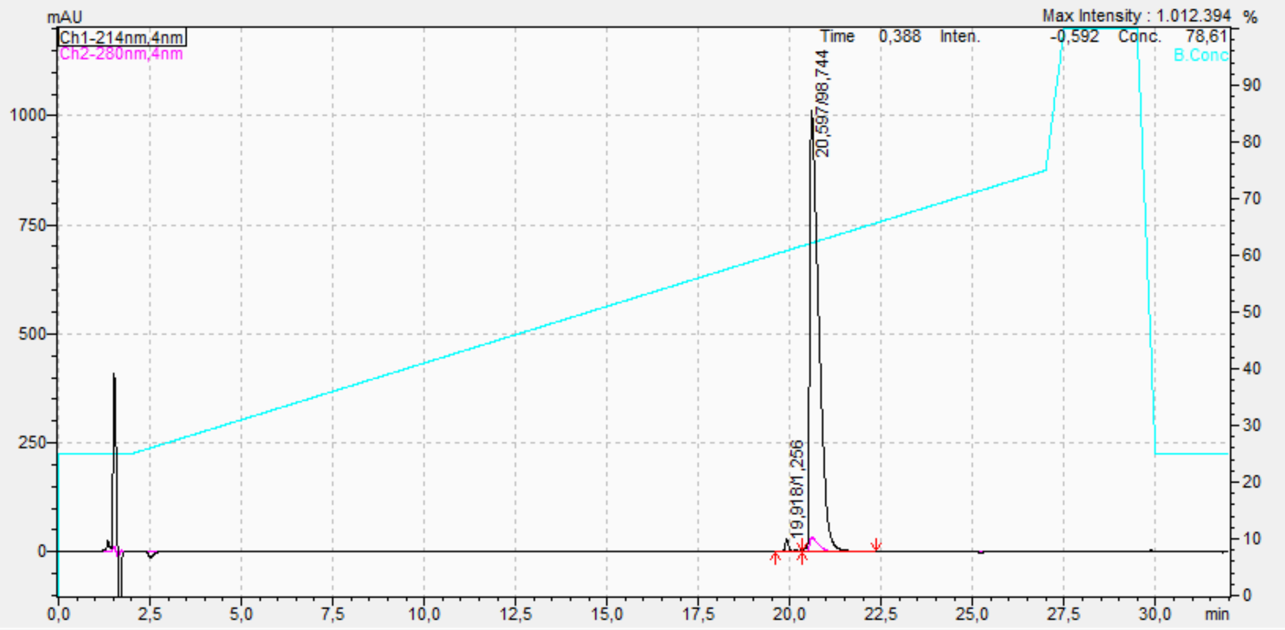

**C12C8 purity.** Analytical HPLC chromatogram.

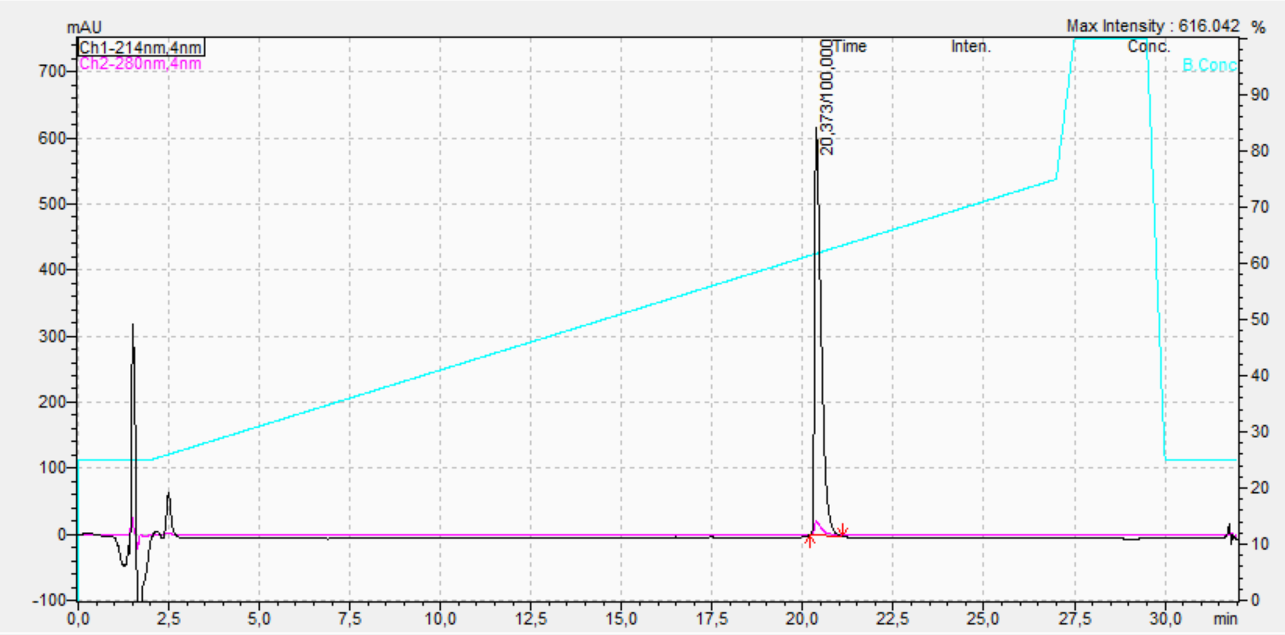

**C8C12 purity.** Analytical HPLC chromatogram.

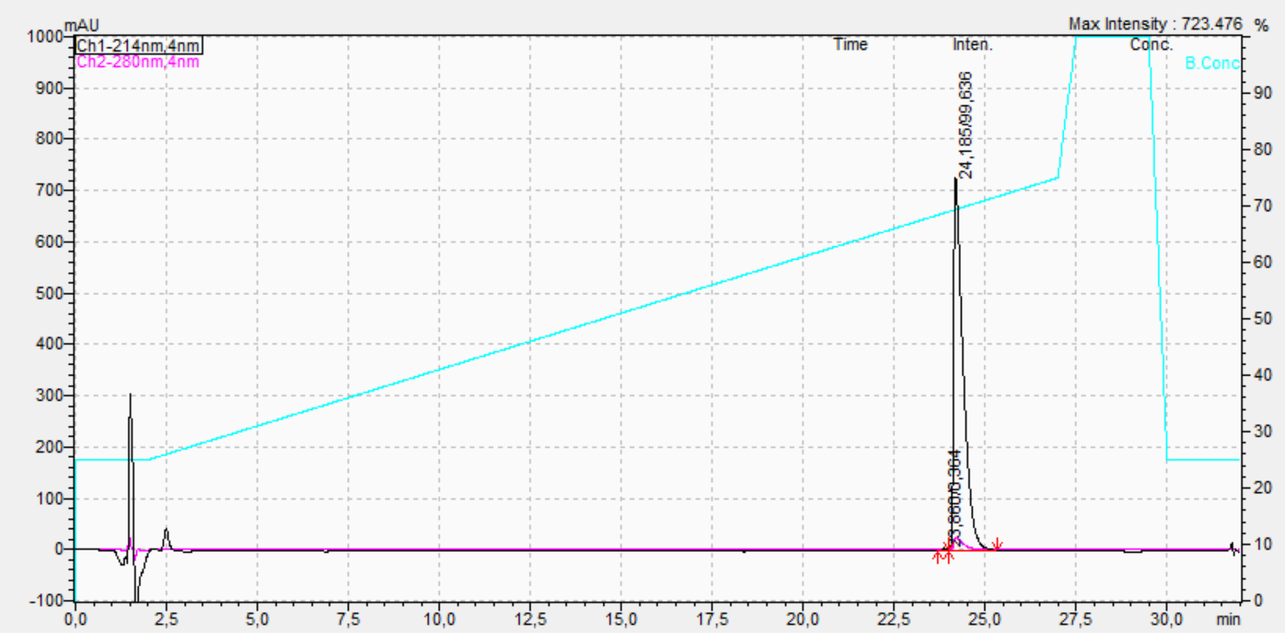

**C12C12 purity.** Analytical HPLC chromatogram.

#### Theoretical calculation of the hydrodynamic radius for a monomeric native salmon calcitonin (sCal) peptide

To estimate a theoretical diameter of a single sCal peptide we applied the theory of determining globular proteins volume found by Harpaz et al. [1]. They outline that the volume of a globular protein can be theoretically determined by applying the partial specific volume of the protein (v) and molecular weight (m) of the protein in the following formula:

$$\frac{v \cdot m}{N_A} = V_{protein}$$

Here $N_A$ is Avogadro constant and $V_{protein}$ is the protein volume. The molecular weight of sCal (CAS number: 47931-85-1) is 3431.85 g/mol. Assuming a monomeric sCal is a globular protein the volume is

$$V_{sCal} = \frac{0.73\ \frac{cm^3}{g} \cdot 10^{21}\frac{nm^3}{cm^3} \cdot 3431.85\frac{g}{mol}}{6.02 \cdot 10^{23}\frac{molecule}{mol}} = \ 4.16\ \frac{nm^3}{molecule}$$

Applying the formula for a sphere we get a hydrodynamic radius of 0.9978 nm for a monomeric native sCal, thus a diameter of 1.995 nm.

#### Brief description of the Nile red assay

IN our study we employed a nile red assay which determines the surface polarity of a hydrophobic aggregate based on the maximum fluorescence signal at 630 nm [2]. Due to the intrinsic hydrophobic property of Nile red, the fluorophore binds to hydrophobic surface regions of an aggregate, resulting in an enhanced fluorescence intensity signal. This method is repeatedly used to characterize the hydrophobicity of various entities such as insulin fibrillation [3], fatty acid containing micelle particles [4] and amyloid fibrillation [5].

### Python based script for semi-automated tracking and analysis of GUVs

To extract kinetic data of reporter dye filling into giant unilamellar vesicles (GUVs) an in-house produced python script was applied. The script consists of three processes: 1) image stack preprocessing, 2) GUV feature detection and temporal linking, and 3) intensity extraction (Fig S1A). The designed script is capable of handling multi-channel images, containing e.g. one GUV membrane channel and at least one reporter dye channel. Throughout the script, Tifffile [6] is applied for image handling, Scikit-image (skimage) [7] and OpenCV are used for image treatment, Numpy [8], Pandas [9] and SciPy [10] are employed for numerical operations and Matplotlib[11] is used for plotting. Below is a detailed description of each of the individual elements in the image analysis.

#### Image stack preprocessing

The GUV recordings were preprocessed by splitting them into individual color channels before flattening them to avoid potential artifacts from a non-evenly distributed illumination profile. The illumination profile induced an almost 40% drop in intensity from center to edge for the reporter dye channels. Flattening was implemented by fitting a second degree polynomial to the full field of view (FOV) (Fig S1B). To ensure that the GUV features did not affect the polynomial fitting the python package skimage.measure [7] was applied to create a FOV mask of the GUV which was passed to the fitting package, from astropy v5.2.1, providing areas not to take into account while fitting. Hereby, the dark spots created by the GUVs dispersing the reporter dyes did not affect the fit. A FOV flattened both with and without taking GUV masks into account is observed in Fig. S1B, cementing the importance of applying the GUV masks for flattening. The features detected for creating the GUV masks for flattening, were sorted based on size and solidity. The raw image FOV was corrected by dividing it by the inverse second degree polynomial in all X and Y coordinates, thus increasing the intensity signal towards the edges while maintaining the signal towards the center. Each temporal frame of each color channel was individually fitted and flattened. Finally, all time frames for each color channel was saved individually.

#### GUV feature detection and temporal linking

Having preprocessed all color channels, the GUV membranes, recorded in the 640 nm channel, were used for creating a mask allowing us to extract the intensity of the reporter dye inside and in the vicinity of each GUV. The python package skimage.measure [7] was applied to detect the GUV, using the following three acceptance criteria for detected features: 1) pixel areas between 500 and 20.000, 2) mean intensity values below 225 and 3) an eccentricity value above 0.9. The centroid coordinates, frame number, bounding box coordinates, perimeter and eccentricity were registered for all accepted GUVs. The detected features were linked using Trackpy [12], creating temporal GUV traces. A search range of 50 was applied with a memory of two frames allowing for GUV X/Y drift and fluctuation in the Z-axis. At this point, non-unilamellar GUVs and traces of less than 20 frames were removed from the registered data. The remaining accepted GUV traces were assigned a unique GUV trace ID consisting of 10 arbitrary numbers and letters allowing us to identify a specific GUV trace along the data treatment. The final procedure in this step was to create small image stacks of 200 by 200 pixels centered around each accepted GUV trace in all recorded color channels.

#### Intensity extraction and quality assessment

To extract the local background and lumen intensity signal in the reporter dye channel of each trace, an adaptive edge-finding algorithm was implemented. The GUV membrane channel of each small image stack was segmented using watershed from skimage [7] creating ring structures which were filled using fillPoly from OpenCV (Fig S1C). The center GUV mask was selected. These image processes were executed for each time frame in each small image stack, thus creating an adaptive GUV mask. The created GUV mask was shrunken using binary_erosion from SciPy [10] to create a mask allowing extraction of the GUV lumen intensity. By using binary_dilation from SciPy [10] a donut shaped mask was created, allowing for extracting the local background intensity just beyond the GUV membrane (see figure S1D for example masks). The average and sum intensities of both GUV local background and lumen intensities from all reporter dye channels were extracted for each time frame and registered. A final data file contained all accepted GUV data, including GUV centroid coordinates, bound box coordinates, perimeter, eccentricity, unique GUV trace ID, background intensities and lumen intensities for each reporter dye channel. For each GUV trace a video was created with four FOV. The FOV are as displayed in Fig. S1C from left to right: 1) for the original small image stack, 2) feature detection via watershed segmentation, 3) labeling of features detected and 4) overlay of the center mask with original small image stack. These videos were applied in manual quality assessment. Finally, each GUV trace was accessed for quality including removing GUV traces where GUV membranes touch.

#### Supplementary tables and figures

**Table S1. Data from DLS and nile red assay for each sCal variant.**

| sCal variant | Diameter [nm] | Nile red intensity [A.U.] |
|---|---|---|
| **C0C0** | 3.3 ± 0.3 | 80.0 ± 1.4 |
| **C4C4** | 2.1 ± 0.2 | 43.5 ± 9.2 |
| **C4C8** | 3.0 ± 0.1 | 104.0 ± 1.4 |
| **C8C4** | 4.5 ± 0.5 | 304.0 ± 5.7 |
| **C8C8** | 5.6 ± 0.3 | 757.5 ± 118.1 |
| **C4C12** | 5.9 ± 0.7 | 1199.0 ± 198.0 |
| **C12C4** | 5.9 ± 0.3 | 681.5 ± 99.7 |
| **C8C12** | 5.5 ± 0.2 | 1431.5 ± 242.5 |
| **C12C8** | 6.0 ± 0.5 | 905.5 ± 26.2 |
| **C12C12** | 5.7 ± 0.1 | 1451.5 ± 222.7 |

Data is collected from two independent experiments each repeated in triplicate. Values are average ± one standard deviation.

**Table S2. Data from calcein leakage experiment for each sCal variant.**

| sCal variants | Peptide concentration | | | | | | |
|---|---|---|---|---|---|---|---|
| | **0.1 $\mu$M** | **0.3 $\mu$M** | **1 $\mu$M** | **3 $\mu$M** | **10 $\mu$M** | **30 $\mu$M** | **100 $\mu$M** |
| **C0C0** | -2.8 ± 0.6 | -6.0 ± 0.7 | -6.8 ± 1.0 | -7.1 ± 1.5 | -7.3 ± 1.4 | -7.3 ± 1.1 | -6.4 ± 2.0 |
| **C4C4** | -1.8 ± 0.2 | -4.3 ± 0.1 | -6.1 ± 0.7 | -5.3 ± 0.7 | -3.0 ± 0.0 | 2.4 ± 1.9 | 23.7 ± 8.4 |
| **C4C8** | -2.0 ± 1.2 | -2.9 ± 0.5 | 3.3 ± 1.2 | 33.0 ± 19.0 | 66.8 ± 24.8 | 93.7 ± 0.6 | 97.0 ± 1.7 |
| **C8C4** | -1.3 ± 0.4 | -4.5 ± 1.3 | 10.9 ± 4.7 | 54.8 ± 14.3 | 86.6 ± 3.5 | 92.4 ± 2.2 | 101.0 ± 3.5 |
| **C8C8** | 1.8 ± 4.8 | 35.9 ± 15.4 | 76.3 ± 12.2 | 97.5 ± 2.6 | 95.6 ± 3.1 | 94.7 ± 2.7 | 96.5 ± 4.8 |
| **C4C12** | 0.4 ± 1.1 | 8.5 ± 9.9 | 83.0 ± 9.7 | 94.4 ± 1.8 | 93.9 ± 4.1 | 95.3 ± 3.8 | 96.9 ± 7.2 |
| **C12C4** | 3.2 ± 5.2 | 23.7 ± 9.5 | 91.5 ± 2.0 | 93.5 ± 1.5 | 94.5 ± 2.3 | 94.3 ± 2.2 | 92.7 ± 6.4 |
| **C8C12** | 31.8 ± 7.1 | 79.6 ± 17.0 | 98.6 ± 4.7 | 99.1 ± 1.0 | 98.1 ± 0.0 | 95.4 ± 1.5 | 97.1 ± 3.5 |
| **C12C8** | 5.7 ± 1.1 | 58.2 ± 26.9 | 100.1 ± 4.1 | 97.1 ± 3.4 | 97.3 ± 3.0 | 95.1 ± 1.0 | 104.5 ± 0.4 |
| **C12C12** | 4.1 ± 7.5 | 28.1 ± 18.8 | 101.9 ± 4.2 | 95.3 ± 4.9 | 94.4 ± 3.1 | 96.4 ± 1.1 | 88.5 ± 13.4 |

Data is collected from two independent experiments. Values are average ± one standard deviation.

**Table S3. P-values from CVM analysis on the slow rate distributions of calcein influx.**

| | C0C0 | C4C4 | C8C4 | C8C12 |
|---|---|---|---|---|
| **C0C0** | | $2.71 \cdot 10^{-11}$ | $3.31 \cdot 10^{-9}$ | $6.83 \cdot 10^{-11}$ |
| **C4C4** | | | $0.99 \cdot 10^{-3}$ | 0.19 |
| **C8C4** | | | | $0.40 \cdot 10^{-3}$ |
| **C8C12** | | | | |

**Table S4. Statistics of GUV's containing inner GUV's (GiGs). Table shows total amount of recorded GiGs and percent of filled outer and inner GiG chambers for control and each sCal variant.**

| Condition | No. of GiGs | GiG w. filled outer chamber [%] | GiG w. filled inner chamber [%] |
|---|---|---|---|
| Control | 78 | 0.0 ± 0.0 | 0.0 ± 0.0 |
| C0C0 | 95 | 0.0 ± 0.0 | 0.0 ± 0.0 |
| C4C4 | 108 | 52.4 ± 18.1 | 16.3 ± 30.1 |
| C8C4 | 86 | 100.0 ± 0.0 | 100.0 ± 0.0 |
| C8C12 | 50 | 100.0 ± 0.0 | 100.0 ± 0.0 |

Error represents one standard deviation. Data was collected from three independent experiments.

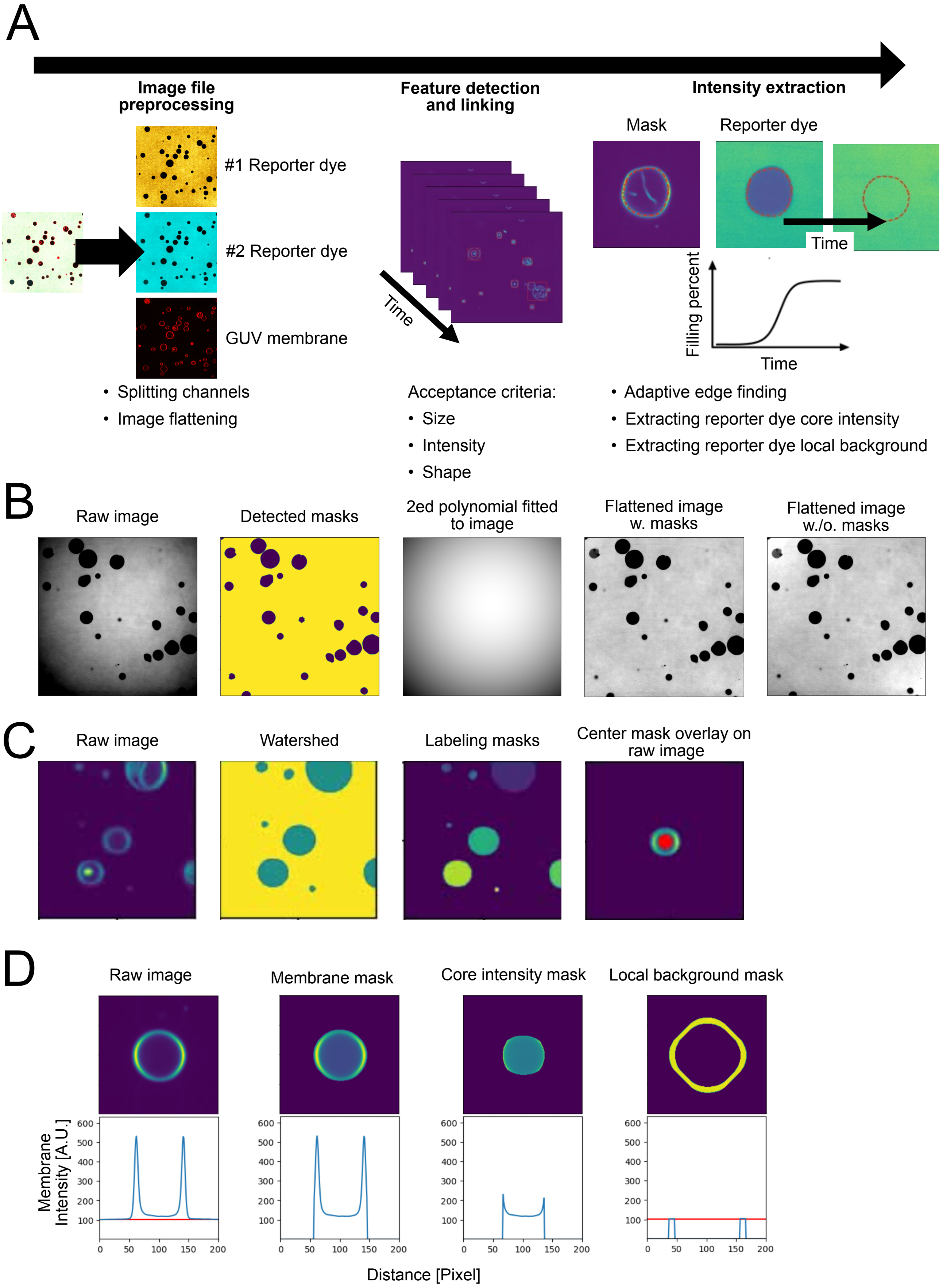
A
Image file preprocessing
#1 Reporter dye
#2 Reporter dye
GUV membrane
• Splitting channels
• Image flattening
Feature detection and linking
Time
Acceptance criteria:
• Size
• Intensity
• Shape
Intensity extraction
Mask
Reporter dye
Time
Filling percent
Time
• Adaptive edge finding
• Extracting reporter dye core intensity
• Extracting reporter dye local background
B
Raw image
Detected masks
2ed polynomial fitted to image
Flattened image w. masks
Flattened image w./o. masks
C
Raw image
Watershed
Labeling masks
Center mask overlay on raw image
D
Raw image
Membrane mask
Core intensity mask
Local background mask
Membrane Intensity [A.U.]
600
500
400
300
200
100
0
50
100
150
200
Distance [Pixel]

**Fig. S1. Flow chart for describing the elements of the semi-automatic GUV tracking and analysis script.** A) Flowchart illustrating the analysis process which consists of 3 steps: image file preprocessing, feature detection, and linking and intensity extraction. B) Images displaying the process of eliminating potential artifacts from an uneven illumination profile by flattening the image. From left to right is displayed 1) raw image, 2) mask of detected features applied for fitting, 3) produced second degree polynomial fit from the data, 4) flattened image using detected GUV masks, and 5) flattened image without using detected GUV masks. C) FOVs in videos centered round detected GUV applied for quality assessment. From left to right is displayed 1) raw image, 2) detected features via watershed segmentation of the raw image, 3) labeling of detected features and 4) an overlay of the center feature used as mask with the raw image. D) Example of applied masks for extracting GUV lumen and local background intensities. Top row displays membrane channel of a single GUV. Bottom row displays the cross section intensity of the image. From left: 1) raw image, 2) raw image overlayed with produced membrane mask, 3) raw image overlayed with produced lumen intensity mask, and 4) raw image overlayed with produced local background mask.

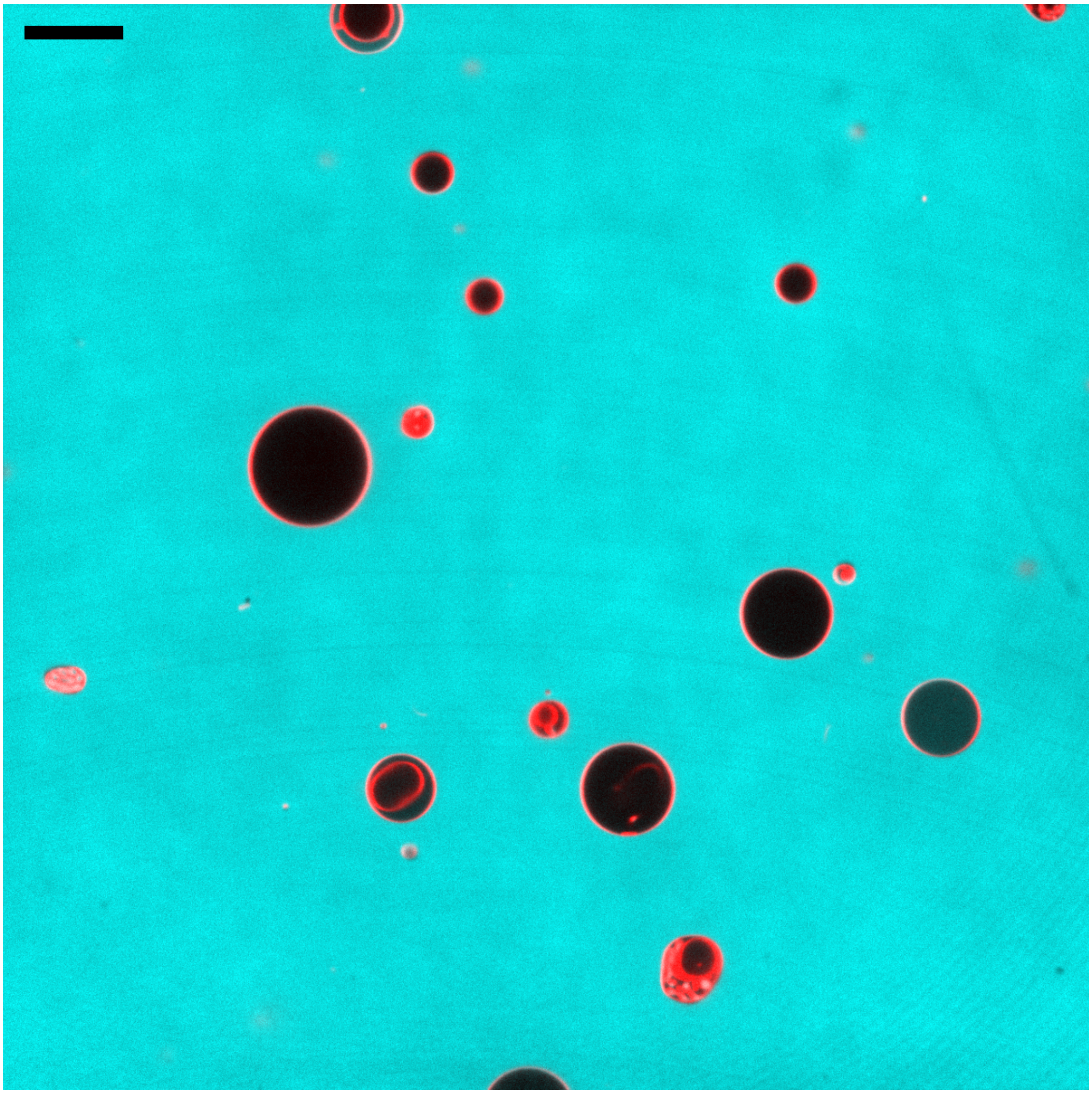

**Fig. S2. Multiple GUVs captured in each recording.** Image displaying typical FOV of a microscopy recording at time point t=0 showing multiple GUVs labelled with DOPE-Atto655 (red) in a calcein solution (cyan). Scale bar represents 20 $\mu$m.

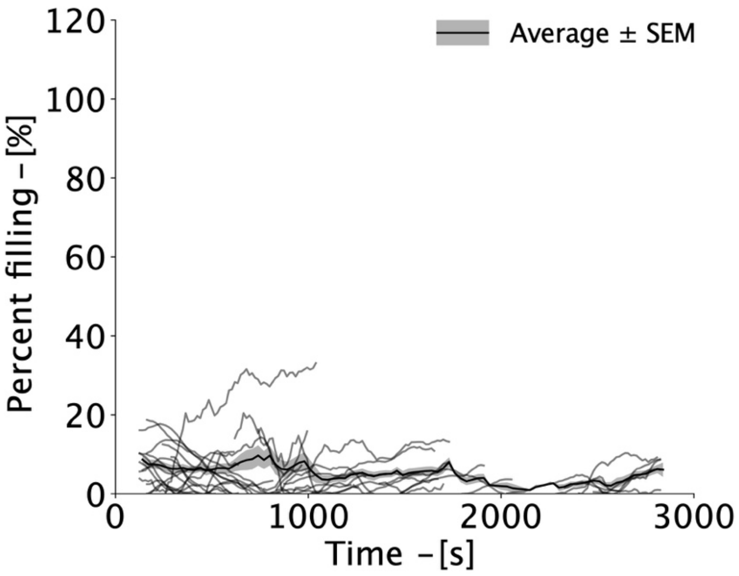

**Fig. S3. Control measurement with no peptide present displays minimal calcein filling.** Percent filling as a function of time for the control measurement with no peptide present. This control show an approximate 10% unspecific leakage of GUVs without any peptides present, which others also have observed [13,14]. N=39 GUVs were collected from two independent recording sessions applying two individual GUV preparations.

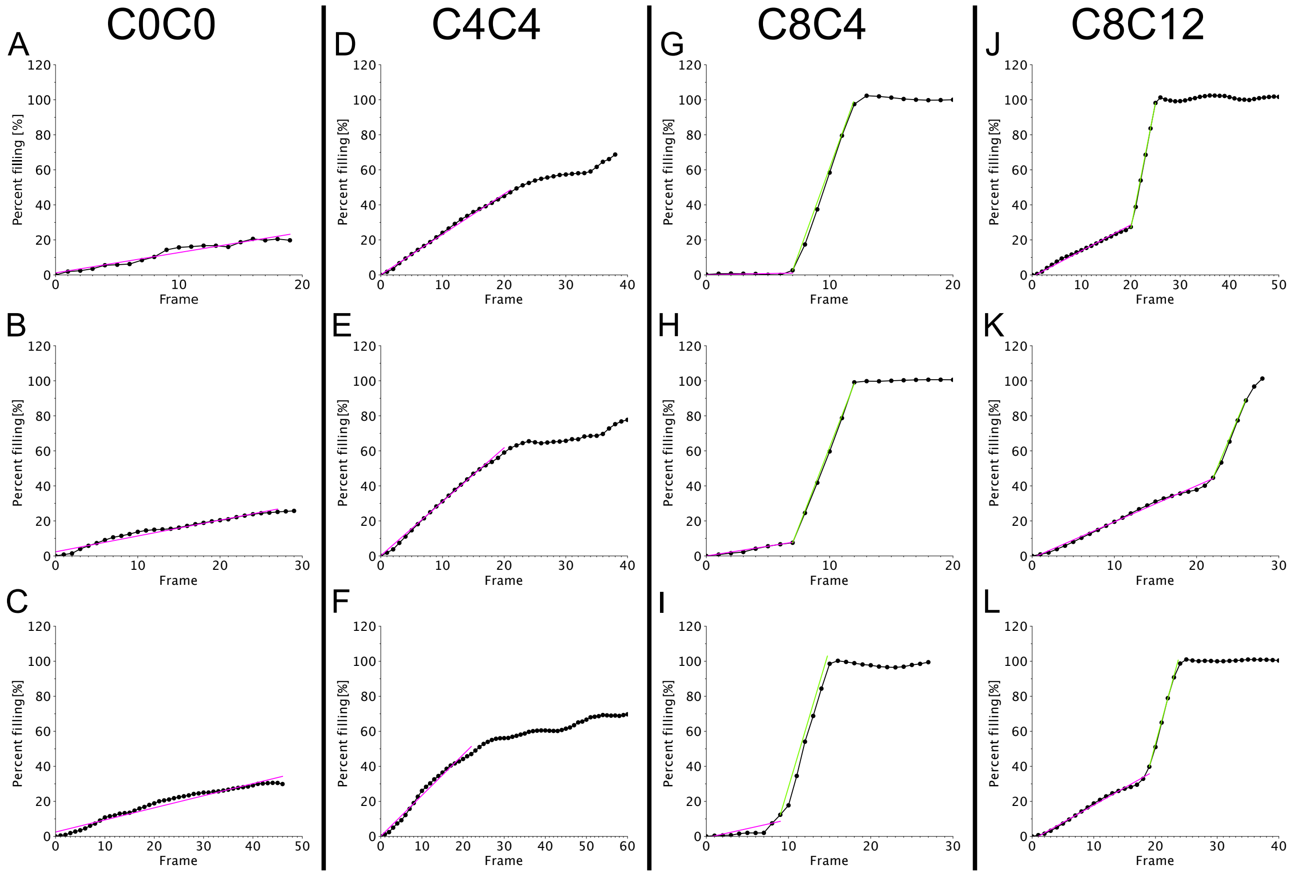

**Fig. S4. Representative calcein filling GUV traces with slow and rapid slope fitting.** Panel of representative fitted GUV traces from each sCal variant: A-C) C0C0, D-F) C4C4, G-I) C8C4 and J-L) C8C12. The magenta line displays regression of points chosen as slow rate while the green line are for rapid rate.

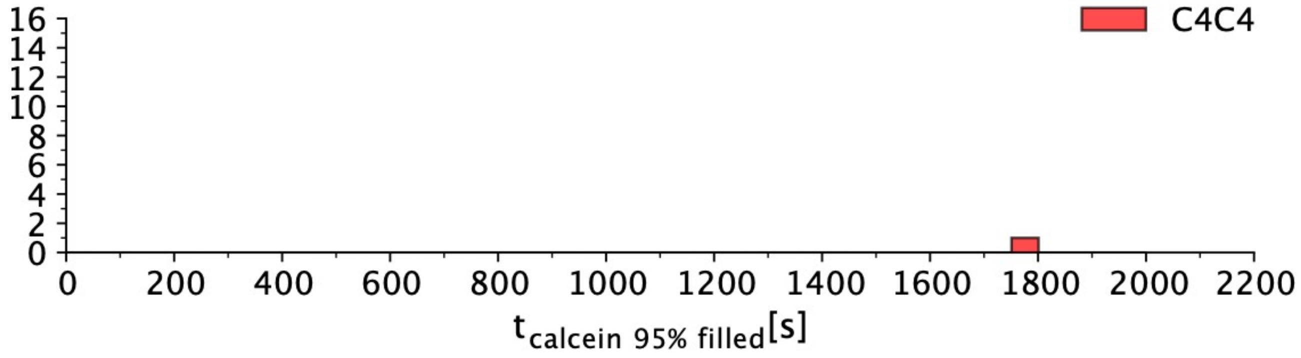

**Fig. S5. Scarce amount of sCal C4C4 traces reach above 95% filling.** Histogram of time spent until a GUV is filled 95% for lipidated sCal variant C4C4 (red), displaying only a single GUV reaching above 95% filling out of 31 GUV traces in total. The single trace was 95% filled after 1772 s.

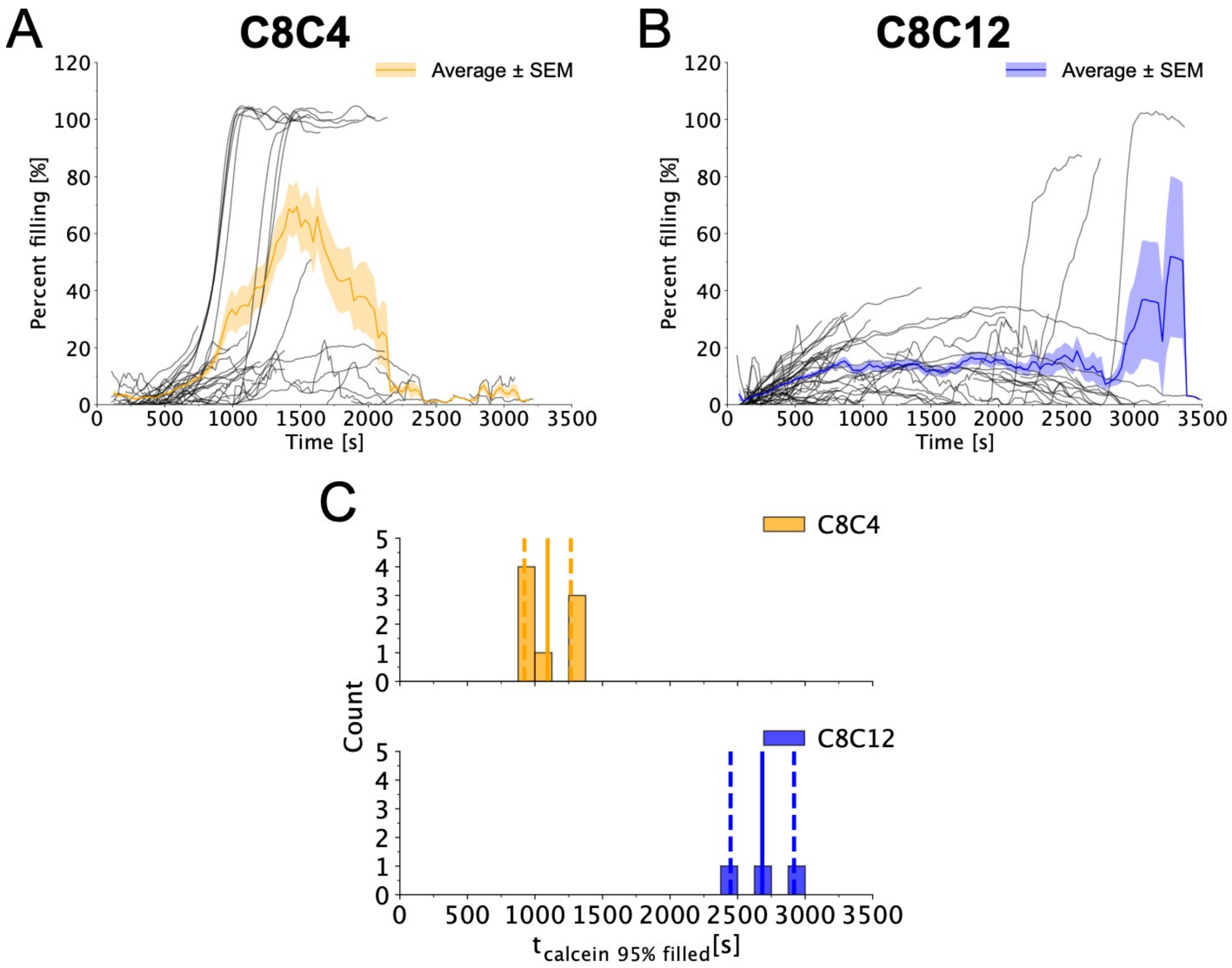

**Fig. S6. C8C4 retained the lower filling time compared to C8C12 at lower peptide concentrations, 30 $\mu$M.** Percent filling of calcein as a function of time for C8C4 (A, yellow) and C8C12 (B, blue), at a peptide concentration of 30 $\mu$M. C) Histogram of time spent until a GUV is filled 95% with calcein ($t_{calcein\ 95\%\ filled}$) for C8C4 (yellow) and C8C12 (blue) at a peptide concentration of 30 $\mu$M. Solid line displays average time spent for the GUVs reaching 95% filled, while dotted lines are one standard deviation, which are 1094 ± 172 s and 2683 ± 235 s for C8C4 and C8C12, respectively. C8C4 induces a significantly faster calcein filling than C8C12 (CVM test, p-value=0.007). Data was collected from three independent recording sessions applying four individual GUV preparations.

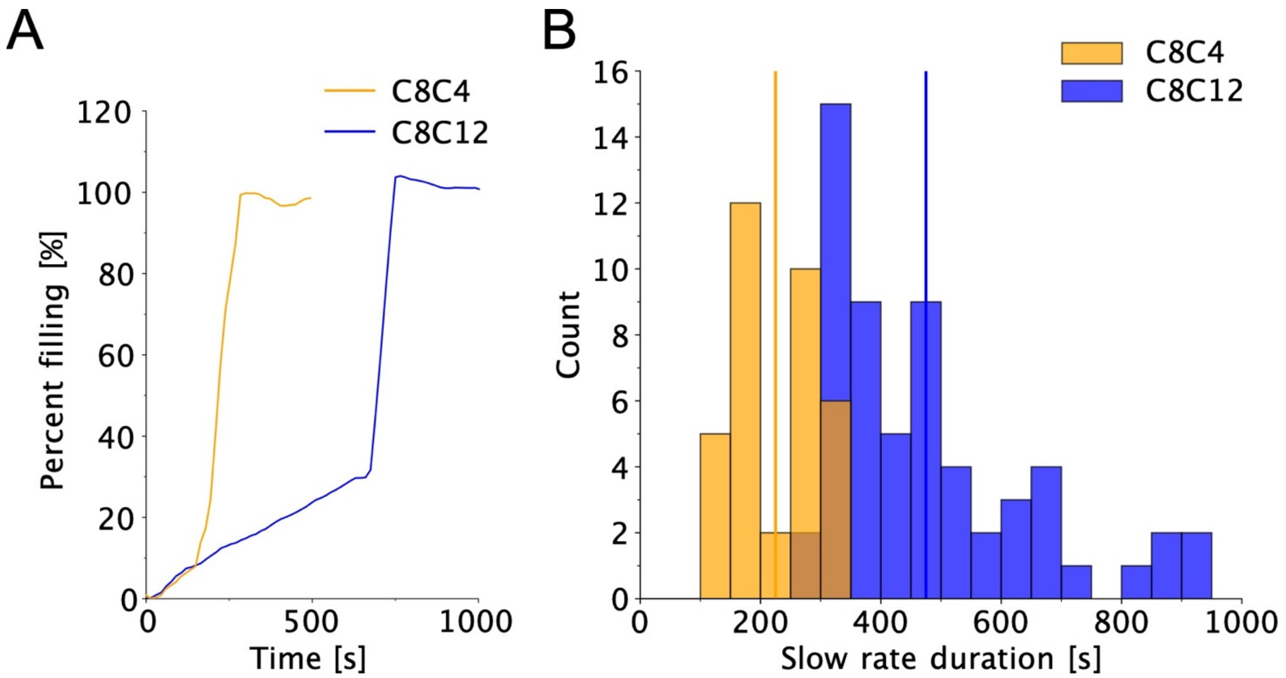

**Fig. S7. C8C4 displays filling profiles with shorter durations in the slow rate regime than C8C12.** A) A representative GUV filling trace from both C8C4 (yellow) and C8C12 (blue) data. The start time of both traces has been set to 0. B) Histogram of slow rate duration, calculated as the time from $t_0$ until the rapid filling rate occurs, for C8C4 (yellow) and C8C12 (blue). Average time is represented with vertical lines. The duration of the slow filling rate phase for C8C4 is significantly shorter than for C8C12 (Cramér-von Mises test, p-value = $6 \cdot 10^{-11}$), with average and one standard deviation values of 225 ± 68 s and 475 ± 167 s, respectively.

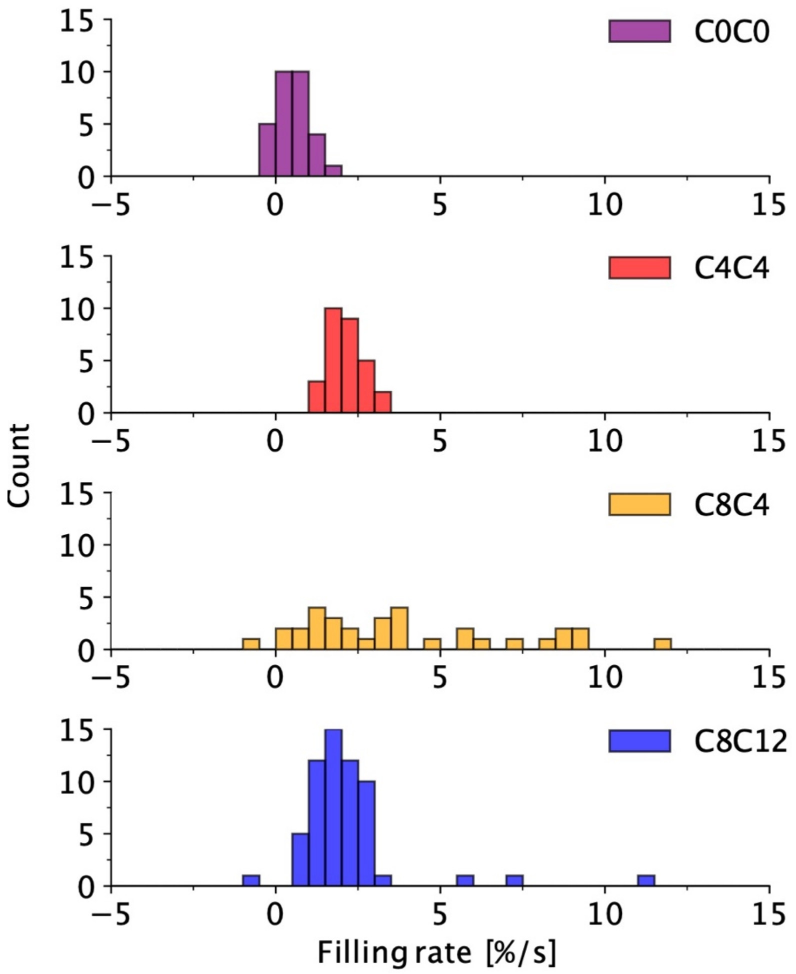

**Fig. S8. Quantified slow calcein filling rates for each selected sCal variant.** Histogram of calcein filling rate values for the slow rates for each GUV trace for lipidated sCal variants, C0C0 (purple), C4C4 (red), C8C4 (yellow) and C8C12 (blue), which obtain an average and one standard deviation of 0.5 ± 0.4 %/s, 2.1 ± 0.5 %/s, 4.6 ± 4.2 %/s and 2.2 ± 1.6 %/s, respectively.

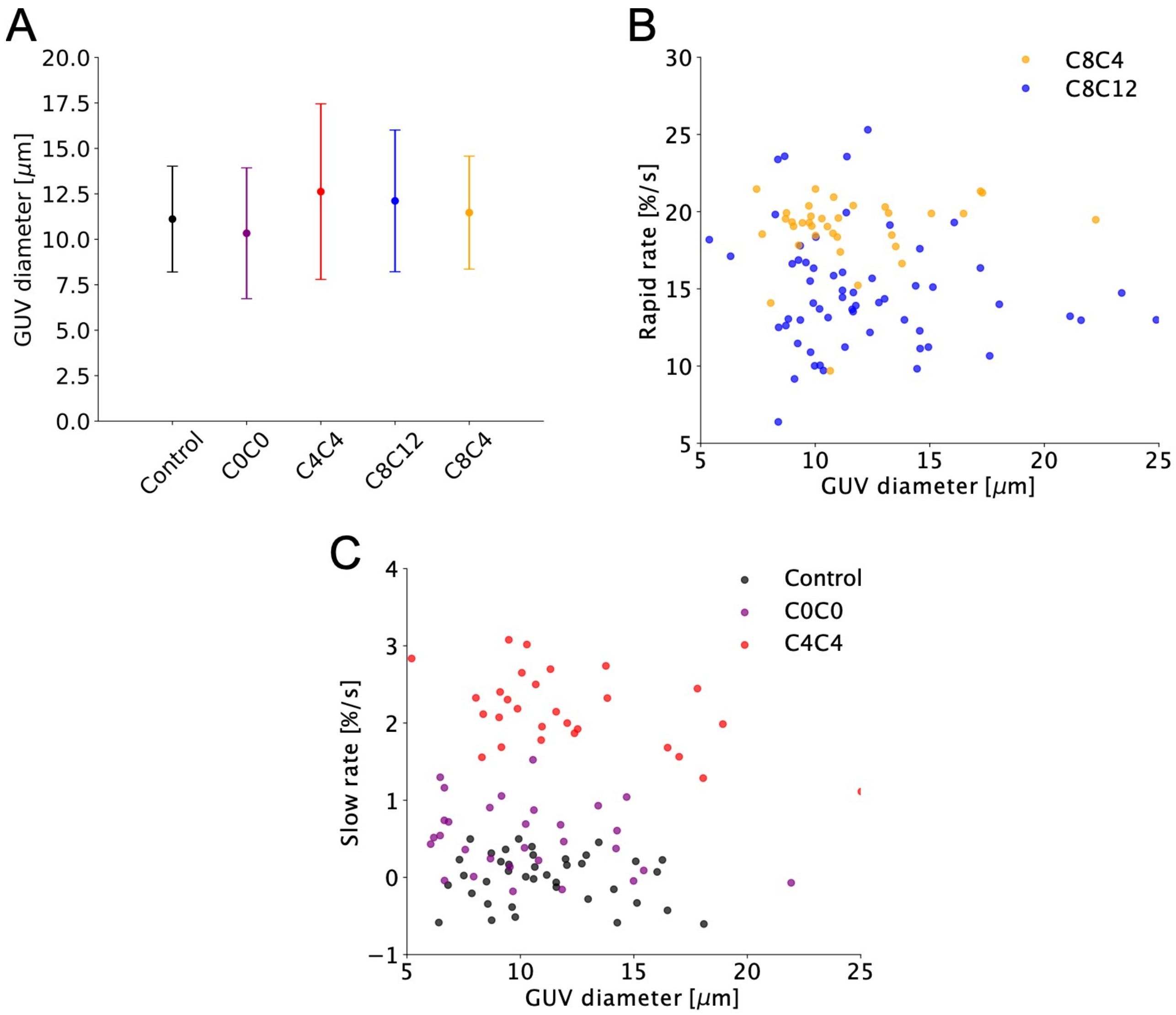

**Fig. S9. No systematic bias introduced by GUV diameter variation.** A) Plot of average size of the GUV population for each of the selected sCal variants and control. Error bars represent one standard deviation. B) Scatter plot of calcein rapid rate filling as a function of GUV diameter for sCal variants C8C4 (yellow) and C8C12 (blue), displaying no systematic dependence of rapid rate as a function of GUV size. C) B) Scatter plot of calcein slow rate filling as a function of GUV diameter for control (black) and sCal variants C0C0 (purple) and C4C4 (red), displaying no systematic dependence of rapid rate as a function of GUV size.

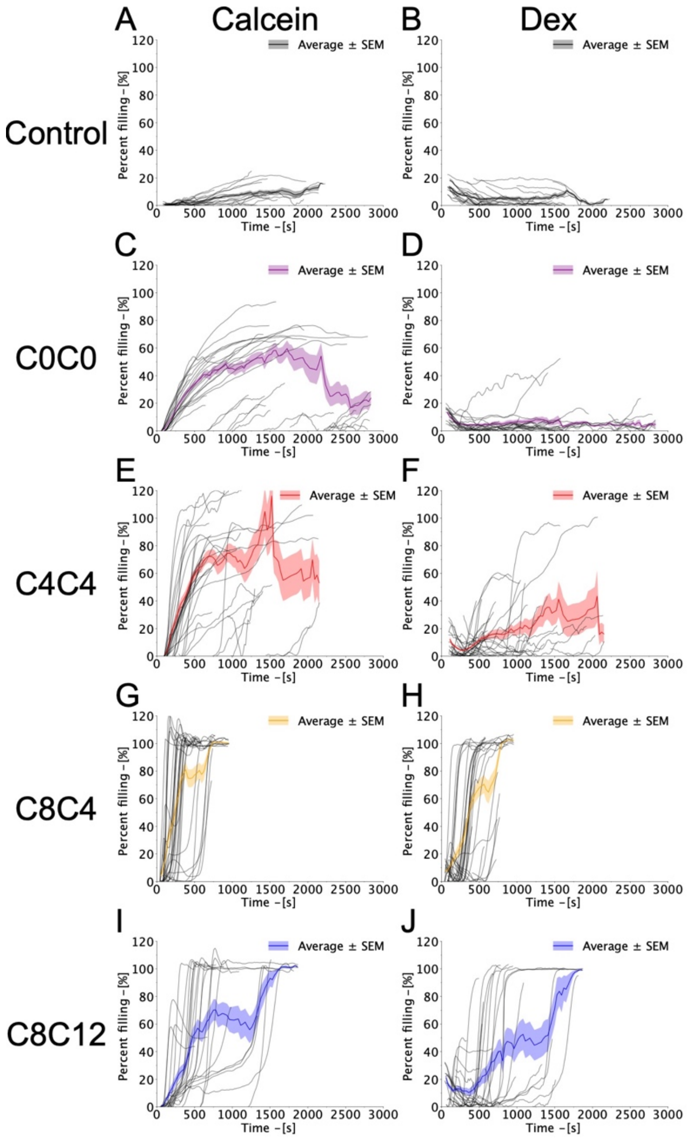

**Fig. S10. Percent filling GUV traces for measurements with both calcein and Dex present.** Percent calcein  and Dex filling as a function of time for the control (A, B) and each sCal variant, C0C0 (C, D), C4C4 (E, F), C8C4 (G, H), C8C12 (I, J), with the gray lines representing the percent filled quantification for each individual GUV trace and the colored line representing the mean of all GUV traces. Plots A, C, E, G, I present influx of calcein in each individual GUV while B, D, F, H, J present Dex influx. The thickness of the colored line represents the standard error of the mean. Data were collected from two independent recording sessions applying two individual GUV preparations.

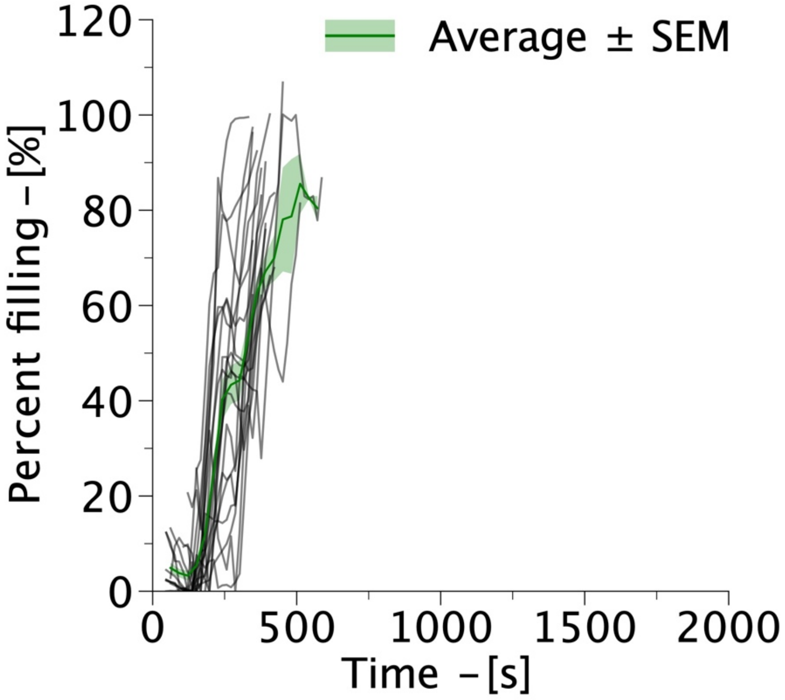

**Fig. S11. Macrolittin-70, a known pore forming peptide, displays rapid rate kinetics.** Percent filling as a function of time for single GUV experiments with 50 $\mu$M macrolittin-70 perturbing POPC GUVs. Data was collected from two independent recording sessions applying two individual GUV preparations.

[8] C.R. Harris, K.J. Millman, S.J. van der Walt, R. Gommers, P. Virtanen, D. Cournapeau, E. Wieser, J. Taylor, S. Berg, N.J. Smith, R. Kern, M. Picus, S. Hoyer, M.H. van Kerkwijk, M. Brett, A. Haldane, J.F. del Río, M. Wiebe, P. Peterson, P. Gérard-Marchant, K. Sheppard, T. Reddy, W. Weckesser, H. Abbasi, C. Gohlke, T.E. Oliphant, Array programming with NumPy,

Nat. 2020 5857825. 585 (2020) 357–362. https://doi.org/10.1038/s41586-020-2649-2.

[9] T. pandas development team, pandas-dev/pandas: Pandas, (2022). https://doi.org/10.5281/ZENODO.7344967.

[10] P. Virtanen, R. Gommers, T.E. Oliphant, M. Haberland, T. Reddy, D. Cournapeau, E. Burovski, P. Peterson, W. Weckesser, J. Bright, S.J. van der Walt, M. Brett, J. Wilson, K.J. Millman, N. Mayorov, A.R.J. Nelson, E. Jones, R. Kern, E. Larson, C.J. Carey, İ. Polat, Y. Feng, E.W. Moore, J. VanderPlas, D. Laxalde, J. Perktold, R. Cimrman, I. Henriksen, E.A. Quintero, C.R. Harris, A.M. Archibald, A.H. Ribeiro, F. Pedregosa, P. van Mulbregt, A. Vijaykumar, A. Pietro Bardelli, A. Rothberg, A. Hilboll, A. Kloeckner, A. Scopatz, A. Lee, A. Rokem, C.N. Woods, C. Fulton, C. Masson, C. Häggström, C. Fitzgerald, D.A. Nicholson, D.R. Hagen, D. V Pasechnik, E. Olivetti, E. Martin, E. Wieser, F. Silva, F. Lenders, F. Wilhelm, G. Young, G.A. Price, G.-L. Ingold, G.E. Allen, G.R. Lee, H. Audren, I. Probst, J.P. Dietrich, J. Silterra, J.T. Webber, J. Slavič, J. Nothman, J. Buchner, J. Kulick, J.L. Schönberger, J.V. de Miranda Cardoso, J. Reimer, J. Harrington, J.L.C. Rodríguez, J. Nunez-Iglesias, J. Kuczynski, K. Tritz, M. Thoma, M. Newville, M. Kümmerer, M. Bolingbroke, M. Tartre, M. Pak, N.J. Smith, N. Nowaczyk, N. Shebanov, O. Pavlyk, P.A. Brodtkorb, P. Lee, R.T. McGibbon, R. Feldbauer, S. Lewis, S. Tygier, S. Sievert, S. Vigna, S. Peterson, S. More, T. Pudlik, T. Oshima, T.J. Pingel, T.P. Robitaille, T. Spura, T.R. Jones, T. Cera, T. Leslie, T. Zito, T. Krauss, U. Upadhyay, Y.O. Halchenko, Y. Vázquez-Baeza, S. 1. . Contributors, SciPy 1.0: fundamental algorithms for scientific computing in Python, Nat. Methods. 17 (2020) 261–272. https://doi.org/10.1038/s41592-019-0686-2.

[11] J.D. Hunter, Matplotlib: A 2D graphics environment, Comput. Sci. \& Eng. 9 (2007) 90–95. https://doi.org/10.1109/MCSE.2007.55.

[12] D.B. Allan, T. Caswell, N.C. Keim, C.M. van der Wel, R.W. Verweij, Trackpy v0.5.0, (2021). https://doi.org/10.5281/ZENODO.4682814.

[13] B. Apellániz, J.L. Nieva, P. Schwille, A.J. García-Sáez, All-or-None versus Graded: Single-Vesicle Analysis Reveals Lipid Composition Effects on Membrane Permeabilization, Biophys. J. 99 (2010) 3619–3628. https://doi.org/10.1016/j.bpj.2010.09.027.

[14] S. Braun, Š. Pokorná, R. Šachl, M. Hof, H. Heerklotz, M. Hoernke, Biomembrane Permeabilization: Statistics of Individual Leakage Events Harmonize the Interpretation of Vesicle Leakage, ACS Nano. 12 (2018) 813–819. https://doi.org/10.1021/ACSNANO.7B08184/ASSET/IMAGES/NN-2017-08184N_M005.GIF.
